# The CD103-XCR1 axis mediates the recruitment of immunoregulatory dendritic cells after traumatic injury

**DOI:** 10.1101/2022.08.19.504399

**Authors:** Ravi Lokwani, Tran B Ngo, Sabrina DeStefano, Kenneth M Adusei, Minhaj Bhuiyan, Aditya Josyula, Mondreakest Faust, Aaron Lin, Maria Karkanitsa, Parinaz Fathi, Kaitlyn Sadtler

**Affiliations:** Section on Immunoengineering, Biomedical Engineering and Technology Acceleration Center, National Institute of Biomedical Imaging and Bioengineering, National Institutes of Health, Bethesda MD 20814; Unit for Nanoengineering and Microphysiological Systems, National Institute for Biomedical Imaging and Bioengineering, National Institutes of Health, Bethesda MD 20814

## Abstract

During wounding and material implantation there is a disturbance in tissue homeostasis and release of self-antigen, and regulation between tolerance and auto-inflammation in injury is not well understood. Here, we analyzed antigen-presenting cells in biomaterial-treated muscle injury and found that pro-regenerative materials enrich Batf3-dependent CD103^+^XCR1^+^CD301b^+^ dendritic cells associated with cross-presentation and self-tolerance. Muscle trauma was accompanied by CD8^+^ iTregs and expansion of CD103^+^XCR1^+^CD62L^-^ adaptive immune cells. Up-regulation of E-Cadherin (the ligand for CD103) and XCL-1 in injured tissue suggests a mechanism for cell recruitment to trauma. Without cross-presenting cells T cell activation increases, pro-regenerative macrophage polarization decreases, and muscle healing is impaired. These data describe a regulatory communication network through CD103^+^XCR1^+^ immune cells resulting in downstream effects on tissue regeneration.

## INTRODUCTION

The goal of tissue engineering is to replace the function of missing or damaged tissues and organs. This can be accomplished using biomaterial scaffolds that integrate and help regenerate injured tissue, as well as medical device implants made of synthetic materials to replace function or cosmesis. When a biomaterial or medical device is implanted in the human body it alters homeostasis and induces a cascade of immune responses that can either positively lead to scaffold integration and tissue growth or yield immune-mediated pathologies (*1, 2*). Thus, understanding how our immune system interacts with engineered materials is necessary to producing next-generation medical devices that work harmoniously with the immune system. Research in this area has grown over the past decade, resulting in a better understanding of how immune cells may moderate material acceptance or rejection and fibrosis. Type 2 immunity has been described in healing of a variety of tissues (*3–5*). Regulatory T cells have also been described in healing processes and this knowledge has been incorporated in material acceptance strategies to try to inhibit excessive immune activation (*6, 7*). Crosstalk between immune cells including T cells, macrophages, monocytes, and fibroblasts, generates an intricate network of cell signaling that contributes to wound healing, regeneration, device acceptance, or inflammation and fibrosis (*8–12*).

Though much of research has focused on macrophages in tissue remodeling and acceptance of engineered materials, biomaterials can also modulate dendritic cell activation (*13, 14*). In evaluating crosstalk among antigen-presenting cells and T cells, we identified a unique conventional dendritic cell subset (CD301b^+^ cDC1) previously implicated in Th2 immunity induction that was enriched by pro-regenerative materials and inhibited by pro-fibrotic materials (*15, 16*). Our model “pro-regenerative” material was decellularized extracellular matrix (ECM), which has been useful clinically in abdominal wall repair and has shown promise in regenerating of traumatic muscle defects (*17*). cDC1s that express CD103 (an integrin that binds E-Cadherin) and XCR1 (a G-protein coupled chemokine receptor that binds XCL-1) are capable of antigen cross-presentation. This process occurs when extracellular antigens are cross-presented onto MHC-I molecules to activate CD8^+^ T cells, which have been associated with self-tolerance (*18, 19*). Furthermore, in systemic studies of autoinflammatory conditions non-canonical HELIOS^+^ CD8^+^ induced regulatory T cells (iTregs) were found to regulate immune responses against self-antigen (*20, 21*).

Here, we identify a Batf3-dependent CD206^+^CD301b^+^ cDC1 subset along with circulating HELIOS^+^CD8^+^ T cells that follow a volumetric muscle loss injury. Without these trauma-associated dendritic cells (tDCs), macrophages lose F4/80 and CD206 expression, markers associated with tolerance and tissue remodeling (*22–24*), T cell activation increases, HELIOS expression on CD8s decreases, and leads to pathologies including ectopic adipogenesis, muscle fiber necrosis, and calcification. These data suggest a role for cross-presentation capable tDCs in response to engineered materials, and subsequent tissue growth and as potential targets for pro-regenerative immunotherapies.

## RESULTS

To evaluate immune responses to engineered materials in trauma and tissue regeneration we utilized a murine model of volumetric muscle loss. After a 3 mm defect was created in the quadriceps muscle group, the resulting void was backfilled with either a control (saline), hydrated decellularized small intestinal submucosa extracellular matrix (ECM) powder, or polyethylene powder (PE) (**Supplemental Figure 1**). ECM was our model “pro-regenerative”, which has been useful clinically in abdominal wall repair and has shown promise in regenerating of traumatic muscle defects (*17*). PE was our model “pro-fibrotic” material, a polymer used in medical devices whose wear particles can cause excessive inflammation and scar tissue deposition (*25*). Material treatment yielded an increase in cellular infiltrate (over control injury) that peaked at 7 days post- injury (ECMtx 28.3 x10^5^ ± 0.3 x10^5^ per quad) and remained at half of the peak infiltration level out to 42 days post-injury (ECMtx 5.3 x10^5^ ± 3.7 x10^5^ per quad). Saline-treated controls peaked cellular infiltration early, by 3 days post-injury (12.3 x10^5^ ± 2.2 x10^5^ per quad **Supplemental Figure 2a**). Histologically, cellular infiltration into the material area increases with overall increases in cell counts observed during flow cytometric analyses. (**Supplemental Figure 2b**).

### Materials recruit a diverse innate immune compartment with a granulocyte shift from eosinophil-dominant to neutrophil-dominant repertoires

A varied set of innate immune cells was recruited to the injury microenvironment within 7 days post-injury, with the number of days dependent upon the material treatment as shown via dimensionality reduction algorithms on resulting flow cytometry data (**Figure 1a-d**). Through manual gating, we identified granulocytes such as neutrophils (main phenotyping marker: Ly6G^+^), basophils (CD200R3^+^), and eosinophils (Siglec-F^+^), as well as mature macrophages (F4/80 and/or CD68^+^) and immature monocyte-like myeloid cells (Ly6C^+^), dendritic cells (CD11c^+^), and other immune cells (CD45^+^Lin^-^) (**Figure 1a**). When comparing different material treatments, we saw a divergence in the immune repertoire by 7 days post-injury (**Figure 1b**). Through the FlowSOM algorithm we identified different sub-populations of macrophages and dendritic cells and confirmed the identification of granulocyte and monocyte-like cell populations (**Figure 1c-d**).

**FIGURE 1.**
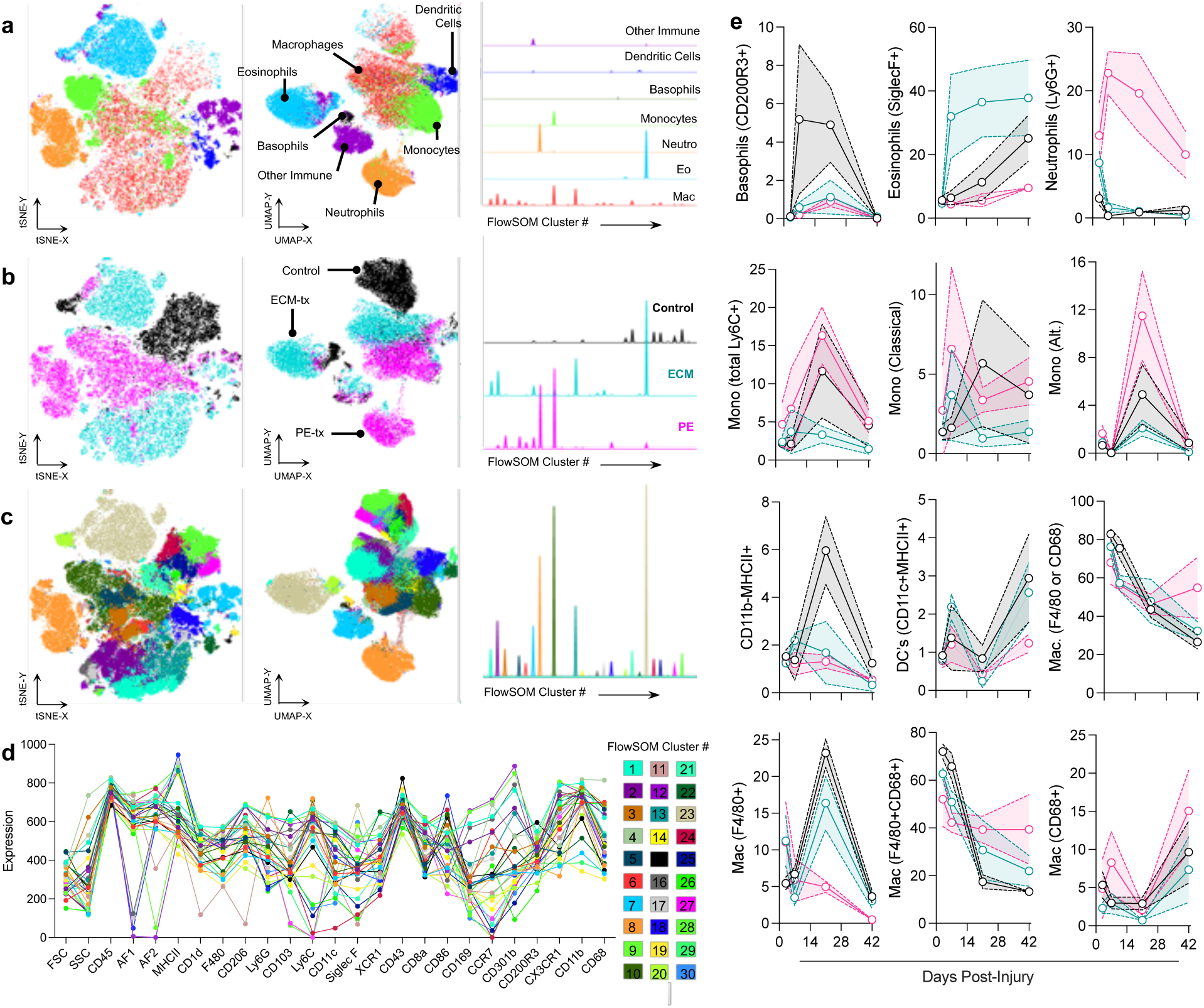
Pro-regenerative and pro-fibrotic materials recruit a diverse range of innate immune cells. Innate immune cell prevalence in biomaterial treated muscle injury. **(a)** t-SNE (left) UMAP (middle) and FlowSOM (right) dimensionality reduction calculations displayed against manually gated populations at 7 days post-injury. Macrophages = red, Dendritic Cells = blue, Monocytes = green, Eosinophils = light blue, Basophils = black, Neutrophils = orange, Other immune cells = purple. **(b)** t-SNE (left) UMAP (middle) and FlowSOM (right) dimensionality reduction calculations displayed against injury treatment type. Control = black, ECMtx = teal, PEtx = pink. **(c)** t-SNE (left) UMAP (middle) and FlowSOM (right) dimensionality reduction calculations displayed against computationally derived clusters (FlowSOM). **(d)** Phenotyping of computationally derived clusters. Expression = scaled fluorescence intensity. **(e)** Immune cell populations over time as a percent of live CD45^+^ immune cells. Basophils: CD11b^-^CD200R3^+^, Eosinophils: CD11b^+^Siglec-F^+^, Neutrophils: CD11b^+^Ly6G^+^, Total Monocytes (CD11b^+^Ly6C^+^), Classical Monocytes Ly6C^hi^, CX3CR1^lo^, Alternative Monocytes: Ly6C^+^CX3CR1^hi^, Non-DC antigen-presenting cells (CD11b^/lo^CD11c^-^MHCII^+^), Dendritic Cells: CD11b^-^CD11c^+^MHCII^+^, Total macrophages: CD11b^+^CD68 and or F4/80^+^, Macrophages: F4/80^+^, Macrophages F4/80^+^CD68^+^, Macrophages: CD68^+^. Control = black, ECM treated = teal, PE treated = pink. Data are means ± 95% Confidence Interval, *n* = 5.

As previously described, we noted a high prevalence of macrophages (F4/80 and/or CD68^+^) that persisted throughout injury recovery (67.9 – 82.8% at 7 days, 54.9 – 26.6 % at 42 days, **Figure 1e, Supplemental Figure 3**) and preferential recruitment of neutrophils (Ly6G^+^) to pro-fibrotic PE-treated (PEtx) muscle injury (22.78% ± 2.72 at 7 days) (*26, 27*). Pro-regenerative materials produced an eosinophil-dominant granulocytic compartment (>30% of CD45^+^ immune cells). CD200R3^+^ Basophils were preferentially recruited to untreated control injuries and peaked between 7 to 21 days post-injury, whereas eosinophils in ECM-treated (ECMtx) injury persisted from 7 through 42 days post-injury. Neutrophils in PEtx peaked by 7 days post-injury and slowly decline by 42 days post-injury while maintaining a large proportion of overall immune cells in the microenvironment. Control and ECMtx injuries both recruited neutrophils early onset, but these cells are cleared 7 days post-injury. PEtx injuries recruit higher levels of Ly6C^hi^ monocytes in comparison to other treatments has been previously reported, with a preference for CX3CR1^+^ cells (11.5% ± 3.0 of cells at 21 days post-injury) (*26*). Dendritic cells were present and persisted in a low proportion (< 4% of total CD45^+^ cells) throughout the time course of response to injury and material implantation. Macrophages peaked early and began to decrease in proportion with time.

### CD103^+^XCR1^+^ dendritic cells are recruited to injury and enriched by pro-regenerative scaffolds

Previous studies have implicated both innate and adaptive immunity in the biomaterial integration and regeneration/fibrosis processes, so we sought to interrogate further the phenotype of antigen-presenting cells (*10, 22, 28*). The majority of MHCII^+^CD45^+^ immune cells in the wound microenvironment are F4/80^+^ macrophages (**Supplemental Figure 4**). Here we found that macrophages in ECMtx muscle injury had higher levels of CD206, CD301b, and CD169 expression suggesting a contribution of the local proliferation of tissue-resident cells with a type- 2 polarization -- a hallmark of Th2-driven inflammation (**Supplemental Figure 5**) (*28*).

While only representing 1 – 2% of the total CD45^+^ immune cell infiltrate, the second most common APCs in the wound space were dendritic cells (CD11c^+^CD11b^lo/neg^). Identification of cross-presentation capable dendritic cells was determined by the expression of XCR1, a GPCR and chemokine receptor, and CD103, an integrin that binds E-Cadherin (**Figure 2, Supplemental Figure 5,6**). CD11b^+^F4/80^+^ macrophages also expressed low levels of CD103 and XCR1, but significantly less than CD11c^+^CD11b^lo^ dendritic cells (**Supplemental Figure 5, 7**). CD103^+^XCR1^+^ Trauma-associated dendritic cells (tDCs) were enriched by pro-regenerative scaffolds. In contrast, pro-fibrotic scaffolds recruited mainly double negative (CD103^-^XCR1^-^) dendritic cells in a pattern that persisted to 42 days post-injury (70% of DCs, **Figure 2a-b**). tDCs expressed intermediate levels of the co-stimulatory molecule CD86 in contrast to CD103^-^XCR1^-^ cells which showed a bimodal distribution with a sub-population of CD86^hi^ dendritic cells (**Supplemental Figure 8**). Neither population expressed CD8⍺, a marker associated with cross presentation, in the muscle tissue. Double negative and XCR1 single positive cells increased in proportion for all groups with time (**Figure 2b,c**), but CD103^+^ XCR1^+^ conventional dendritic cells were peak by proportion by 3 to 7 days post-injury and by count at 7 days post-injury (**Figure 2b,c**). Very few CD103 single positive dendritic cells were in the wound space throughout injury recovery (**Figure 2b,c**).

**FIGURE 2.**
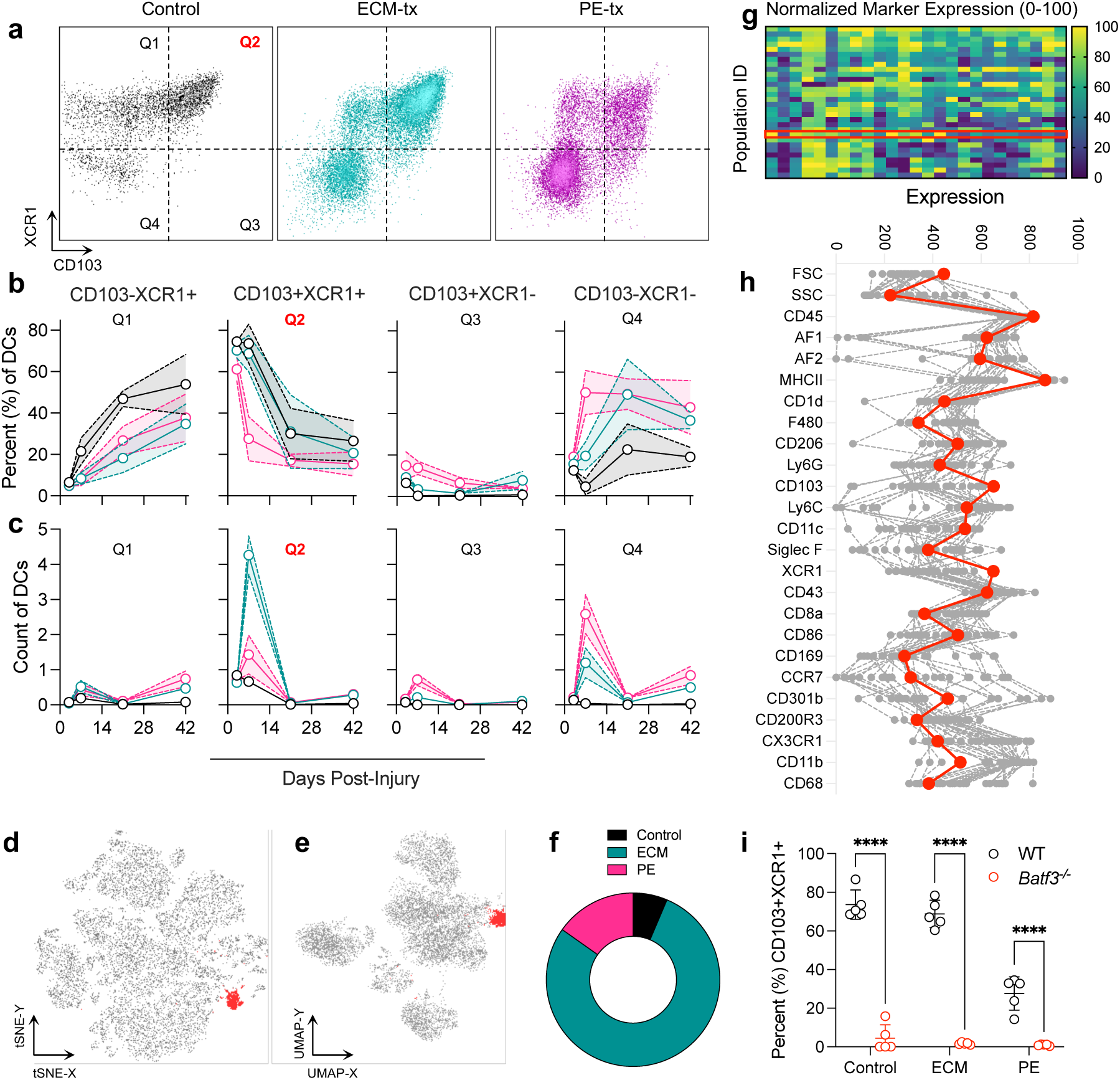
Batf3-Dependent cross-presenting dendritic cells are enriched by pro-regenerative scaffolds and peak by 7 days post-injury. **(a)** Representative dot plots of CD45^+^CD11b^lo/-^CD11c^+^MHCII^+^ dendritic cells at 7 days post-injury. Four quadrants correspond to: Q1 = CD103^-^XCR1^+^, Q2 = CD103^+^XCR1^+^, Q3 = CD103^+^XCR1^-^, and Q4 = CD103^-^XCR1^-^. **(b)** Proportion (%) of dendritic cells within four quadrants at 3-, 7-, 21-, and 42-days post-injury, Control = black, ECM treated = teal, PE treated = pink. **(c)** Count (in 10,000s) of dendritic cells per quad at 3-, 7-, 21-, and 42-days post-injury. **(d)** tSNE and **(e)** UMAP of CD45^+^ immune cells at 7-days post-injury concatenated from three treatment groups, tDC population highlighted in red. **(f)** Proportion of tDC population from three treatment groups. **(g)** Phenotyping of 30 different immune cell clusters identified by FlowSOM algorithm, normalized within each marker where min = 0, and max = 100. **(h)** FlowSOM scaled expression values on different clusters with cluster 22 (tDCs) highlighted in red. **(i)** Proportion of CD11c^+^MHCII^hi^ dendritic cells that are positive for CD103 and XCR1 at 7 days post-injury in WT (black) and Batf3-deficient (*Batf3^-/-^*) mice. Data are means ± 95% Confidence Interval, *n* = 5 representative of at least two independent experiments.

When evaluating cell populations through dimensionality reduction and hierarchical clustering algorithms, the FlowSOM algorithm identified tDCs as a unique cluster. Cells mapped to an island with both t-SNE and UMAP visualizations (**Figure 2d,e, Supplemental Figure 9**). When FlowSOM was applied to the three treatment groups within the same clustering population, over 70% of the cells within the tDC cluster were from ECMtx injury whereas only 15% and 6% were from PEtx and Control injuries, respectively (**Figure 2f-h**). Identification of these cells was confirmed in two C57BL/6 mouse litters as well as in the muscle injury of a Lewis Rat, suggesting this is repeatable and applicable to multiple species within the Muridae subfamily (**Supplemental Figure 10**) (*29*). To determine whether tDCs were dependent upon BATF3, a transcription factor required for the differentiation of cross-presenting dendritic cells (*30*), we evaluated the myeloid compartment in cross-presentation deficient *Batf3^-/-^*mice. In these mice, there was a loss of CD103^+^XCR1^+^ dendritic cells, suggesting tDCs are dependent upon the same signaling pathways as CD8⍺^+^ cross-presenting dendritic cells described in the literature (ECMtx WT 68.78% ± 7.09 versus *Batf3^-/-^* 1.65% ± 0.56, *P* = 1.74 x 10^-14^, **Figure 2i**). At 7 days post-injury when tDCs peaked in the muscle tissue, they were also detected in the overlying skin incision site of ECM-treated muscle injury, but by a significantly lower fraction (**Supplemental Figure 11**).

Outside of the local muscle tissue, dendritic cells were evaluated in the blood and lymph nodes of injured mice. These DCs (CD11b^-/lo^CD11c^+^MHCII^hi^) had a differing expression of markers associated with antigen cross-presentation capacity (XCR1, CD103, and CD8⍺) depending upon their location (**Supplemental Figure 12**). In muscle, tDCs expressed high levels of both XCR1 and CD103, with no expression of CD8⍺. In the lymph node, these cells expressed all three markers, XCR1, CD103 and CD8⍺, whereas in the blood most DCs were XCR1^-^CD103^-^CD8⍺^-^. Interestingly, there was a larger proportion of CD11c^+^MHCII^hi^ DCs in the blood of PE-treated mice. Most of these cells (> 40%) were B220/CD45R^+^ suggesting a potential preference towards circulating pDC recruitment for pro-fibrotic material implants (**Supplemental Figure 13a-c**). In the lymph node there was no significant difference in the proportion of dendritic cells, though PE-treated mice did have a higher proportion of TCRɣδ^+^ T cells agreeing with previous literature on the role of these cells in fibrotic disease (**Supplemental Figure 13a**).

**FIGURE 3.**
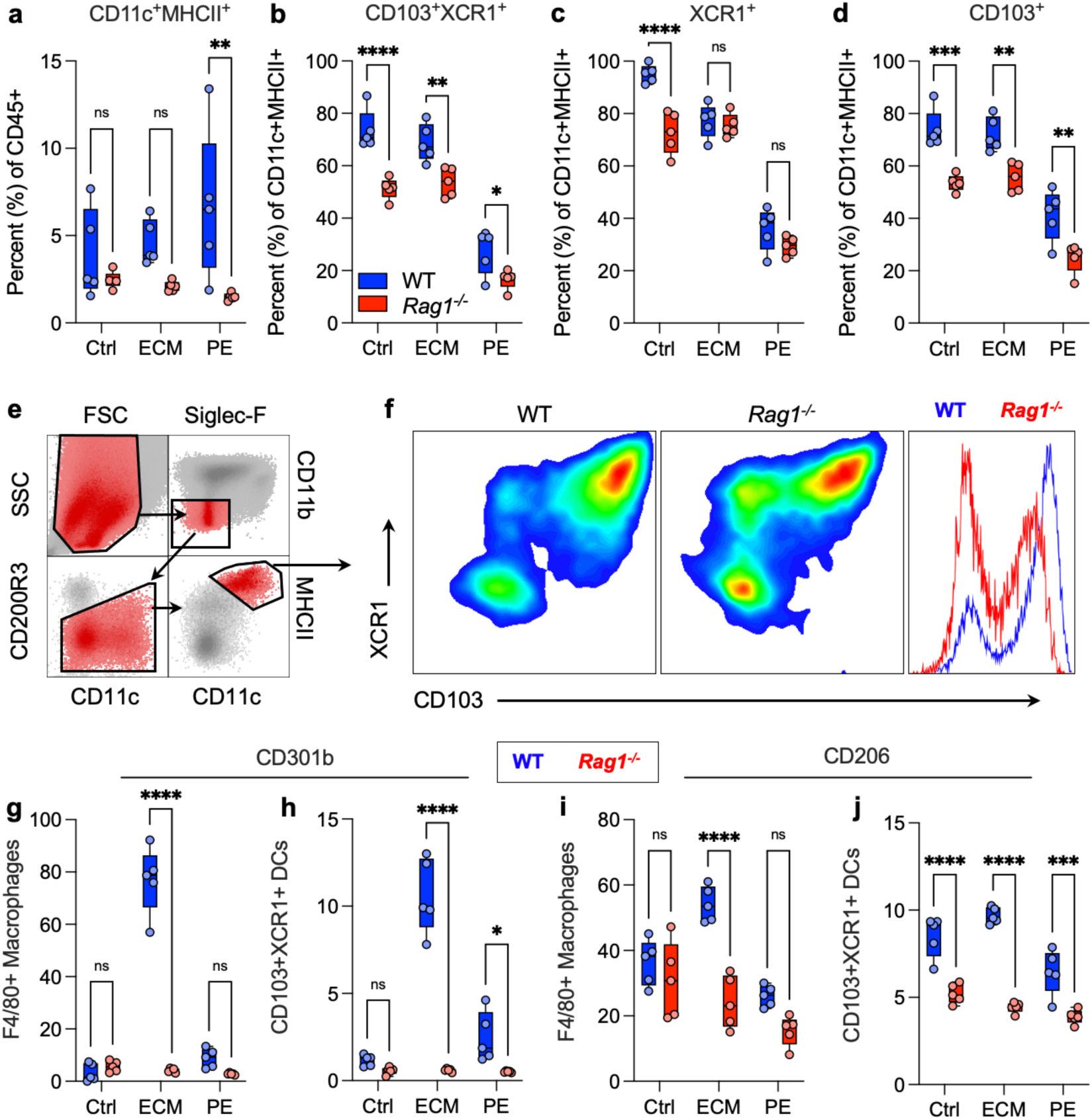
Type-2 myeloid markers are enhanced on cross-presenting dendritic cells by pro- regenerative scaffold treatment and depend on adaptive immunity. **(a)** CD11c prevalence in muscle tissue as a proportion of CD45^+^ live immune cells **(b)** CD103^+^XCR1^+^ dendritic cells as proportion of total DCs **(c)** CD103^+/-^XCR1^+^ dendritic cells. **(d)** CD103^+^ XCR1^+/-^ dendritic cells. **(e)** Gating strategy for dendritic cells **(f)** Representative FACS plots. Blue = Wild Type C57BL/6J mice, Red = *Rag1^-/-^* B6 mice. **(g)** CD301b expression on F4/80^+^ macrophages in wild type (WT) and RAG-deficient mice (*Rag1^-/-^*). **(h)** CD301b expression on tDCs in WT and *Rag1^-/-^* mice. **(i)** CD206 expression on F4/80^+^ macrophages In WT and *Rag1^-/-^* mice. **(j)** CD206 expression on tDCs in WT and *Rag1^-/-^* mice. ANOVA with Tukey Post-hoc correction for multiple comparisons, * = *P* < 0.05, ** = *P* < 0.01, *** = *P* < 0.001, **** = *P* < 0.0001, *n* = 5.

### Phenotype and recruitment of CD103^+^XCR1^+^ dendritic cells and scaffold-associated macrophages are dependent on adaptive immunity

As dendritic cells directly communicate with adaptive immune cells such as T cells and B cells through antigen presentation, we sought to evaluate the inverse relationship and role of adaptive immunity on recruitment and activation of dendritic cells within the scaffold microenvironment. In RAG-deficient mice (*Rag1^-/-^*), which lack T cells and B cells, there were similar proportions of dendritic cells recruited to the overall environment, trending fewer in *Rag1^-/-^*, but significantly fewer XCR1^+^CD103^+^ tDCs as a proportion of total dendritic cells (**Figure 3a-f**). When evaluating tDC recruitment based on expression of XCR1 or CD103, the partial loss of CD103 expression was the main contributor to the loss of the tDCs, given that XCR1 expression was largely maintained in the absence of adaptive immune cells. As with macrophages, tDCs in wild type mice had high levels of CD301b and CD206 in the presence of ECM scaffolds (**Figure 3g,h**). ECM scaffold-mediated type-2 upregulation was lost for both CD301b and CD206 (**Figure 3g-j**) in *Rag1^-/-^* mice when compared to wild type mice.

### Recruitment of NK cells and prevalence of HELIOS^+^ iTregs in local tissue and periphery after trauma

At 7 days post-injury B cells, NK cells, ⍺β T cells, and γδ T cells were detected in the injury microenvironment (**Figure 4a**). ECMtx up-regulated the number and proportion of CD49b^+^ NK cells in comparison to both control injury and PEtx injury (2.498% ± 1.224 versus 0.216% ± 0.082 and 0.0734% ± 0.017, *P* < 0.0001). Regarding ⍺β conventional T cells, there was a larger proportion of CD4^+^ helper T cells in comparison to CD8^+^ T cells in all treatment groups, though the ECMtx enriched a higher proportion of CD4s compared to control and PEtx (73.08% ± 3.81 versus 57.48 ± 5.17 and 57.18 ± 3.25, **Figure 4b**). The diversity of non-myeloid cell types in the injury microenvironment could be visualized via t-SNE and UMAP of corresponding data and displayed a diverse population of adaptive immune cells (**Figure 4c**). Within the CD4^+^ and CD8^+^ T cell compartments we found a large proportion of cells that expressed FoxP3 (for CD4s) and HELIOS (for CD4s and CD8s). The ECMtx greatly enriched HELIOS^+^ CD8^+^ iTregs compared to PEtx (80.74% ± 11.08 versus 33.14% ± 17.62, *P* < 0.0001). This was also prevalent in the blood and draining lymph node by 21 days post-injury, where HELIOS expression in peripheral blood CD8s was readily apparent (**Figure 4e,f**). Additionally, we found a population of ST2^+^ B Cells which have previously been described as regulatory B Cells, that were ablated in *Batf3^-/-^* mice (ECMtx WT 0.458% ± 0.099 versus 0.25% ± 0.016, *P* = 0.0007, **Supplemental Figure 14**).

**FIGURE 4.**
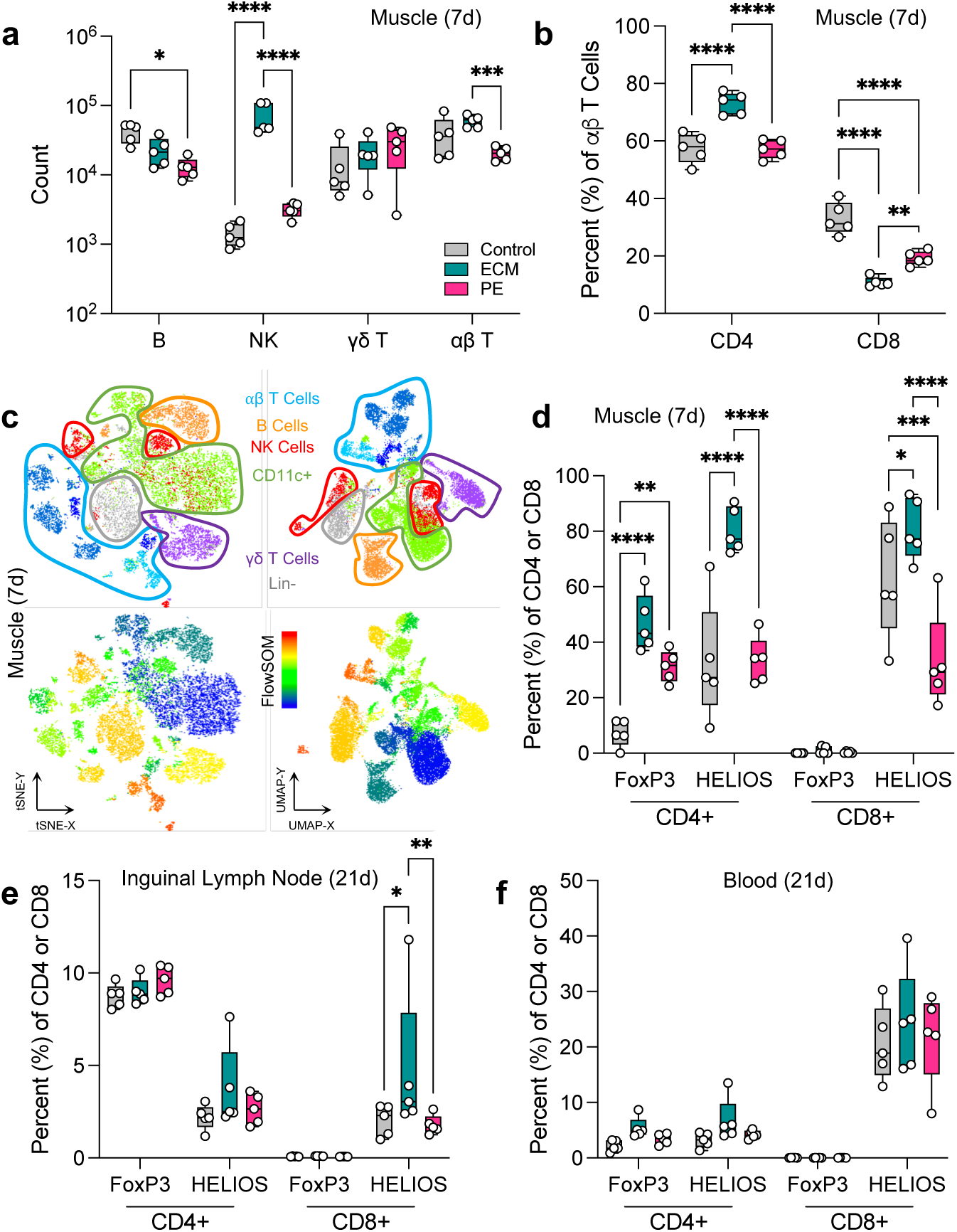
Prevalence of NK Cells and HELIOS+ iTregs enriched by pro-regenerative material treatment. **(a)** Count of adaptive immune cells within the injury space at 7 days post-injury. **(b)** Proportion of ⍺β T cells that are either CD4^+^ or CD8^+^. **(c)** tSNE and UMAP projections of CD11b^-^ compartment at 7 days post-injury. **(d)** Proportion of FoxP3^+^ and HELIOS^+^ cells in the injury space at 7 days post-injury. **(e-f)** Proportion of Tregs in the **(e)** inguinal (draining) lymph node and **(f)** peripheral blood at 21 days post-injury. ANOVA with Tukey Post-hoc correction for multiple comparisons, * = *P* < 0.05, ** = *P* < 0.01, *** = *P* < 0.001, **** = *P* < 0.0001. *n* = 5.

### Loss of cross-presentation results in over-activation of CD4+ T cells, inhibition of pro-regenerative macrophage polarization, and muscle pathology

In addition to these regulatory T cells, we found a sub-population of CD103^+^XCR1^+^ adaptive immune cells induced by trauma by 7 days post-injury that was still present by 21 days post-injury in the draining lymph nodes (**Figure 5a-b, Supplemental Figure 15**). This population was present for B cells, CD4^+^ T cells, CD8^+^ T cells, and ɣδ T Cells and increased with time (**Supplemental Figure 16a,b**). In B Cells and CD4^+^ T cells, most of this population was CD62L^-^ even without injury, and for CD8^+^ and ɣδ T Cells activation increased with time, with CD8s reaching their peak by 7 days post-injury and ɣδ T Cells increasing through 21 days post injury (**Supplemental Figure 16**). In addition to the CD103^+^XCR1^+^ population, there was also a CD103^lo^XCR1^-^ population that was prominent in CD8^+^ T cells and ɣδ T Cells (**Supplemental Figure 16d,e**). Activation of CD103^+^XCR1^+^ adaptive immune cells, as determined by the loss of CD62L expression, peaked by 7 days post injury. They were enriched in the local tissue compared with peripheral blood and the draining lymph node (**Supplemental Figure 17,18**). We found that the proportion of HELIOS^+^ iTregs that were CD103^+^XCR1^+^ increased over time, more so than FoxP3^+^HELIOS^-^ Tregs. This pattern was true for both CD4^+^ iTregs as well as CD8^+^ iTregs (**Supplemental Figure 19**).

**FIGURE 5.**
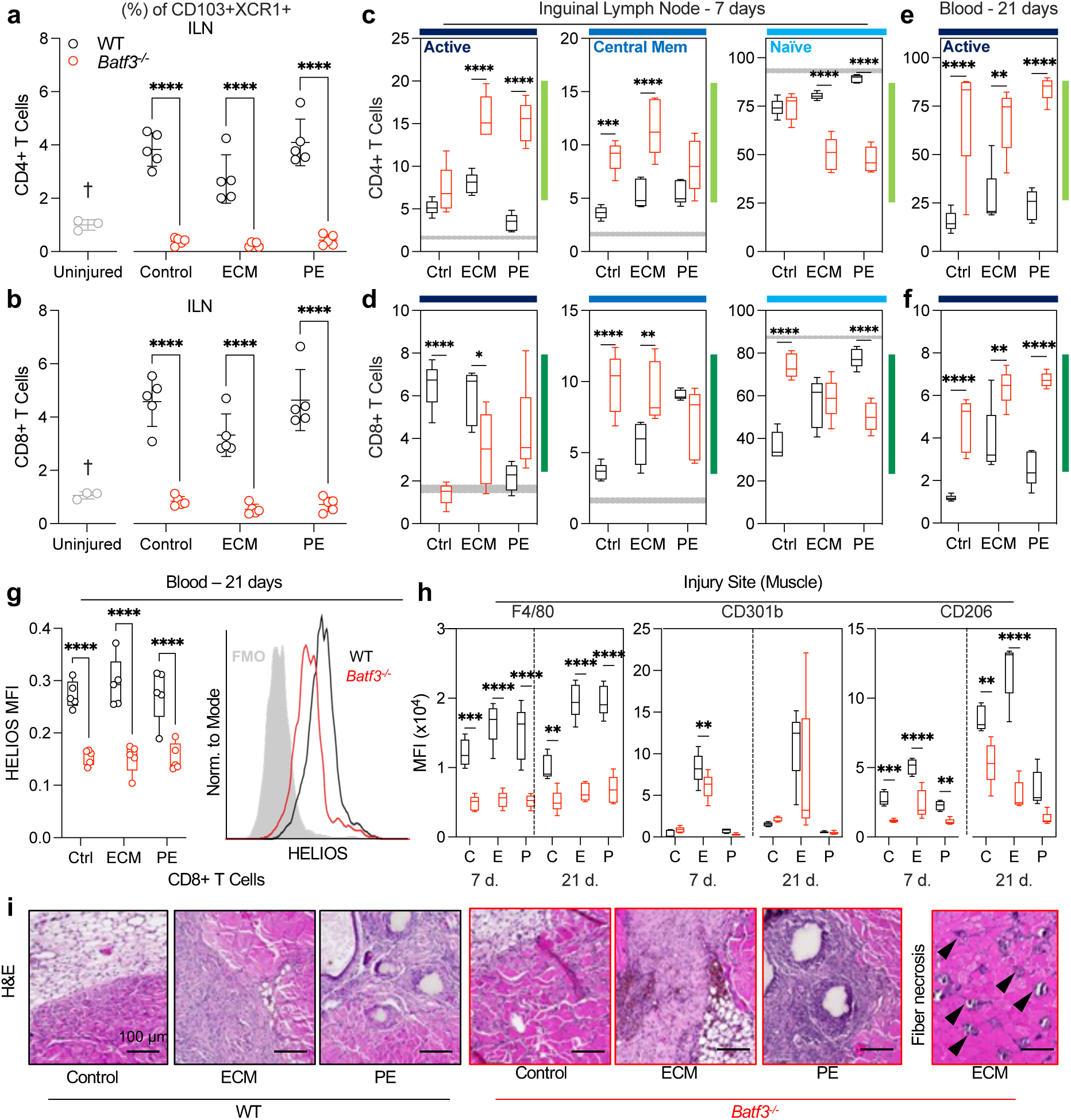
Loss of cross-presenting cells results in over-activation of T cells, decreased pro-regenerative macrophage phenotype, and muscle pathology. (a - b) Proportion of CD4^+^ and CD8^+^ T cells that are CD103^+^XCR1^+^ in the draining lymph node at 7 days post-injury. Grey circles = uninjured control (B6), black = C57BL/6 WT, red = *Batf3^-/-^*. **(c-d)** Activation profile of **(c)** CD4^+^ T cells and **(d)** CD8^+^ T cells in the inguinal lymph nodes of WT (black) and *Batf3^-/-^*mice based on the expression of CD62L and CD44 at 7 days post-injury. Light green = CD4^+^ T cells, Dark green = CD8^+^ T cells. Active/Effector = CD62L^-^ CD44^+^, dark blue; Central Memory = CD62L^+^CD44^+^, medium blue; Naïve = CD62L^+^CD44^-^, aqua. Gray bands in c and d are range in uninjured control lymph nodes (min to max). **(e)** CD4^+^ and **(f)** CD8^+^ T cell activation in peripheral blood at 21 days post-injury. **(g)** HELIOS expression on CD8^+^ T cells in peripheral blood at 21 days post-injury. **(h)** Macrophage polarization at 7- and 21-days post-injury. MFI = mean fluorescence intensity. Ctrl/C = Control, E = ECMtx, P = PEtx. **(i)** Hematoxylin & Eosin (H&E) staining of muscle at 21 days post-injury in WT and *Batf3^-/-^* mice, arrows = fiber necrosis/calcification. Data are means ± SD (a-b) or range (c - g), *n* = 3 - 5, ANOVA with Tukey’s post-hoc correction. * = *P* < 0.05, ** = *P* <0.01, *** = *P* < 0.001, **** = *P* < 0.0001; † = uninjured versus each treatment group *P* < 0.05.

CD103^+^XCR1^+^ adaptive immune cells were absent in the draining lymph nodes of injured cross-presentation deficient *Batf3^-/-^* mice at 7 days post-injury (3 – 5% versus 0.3 – 1%, **Figure 5a-b, Supplemental Figure 20**). In the lymph node of *Batf3^-/-^* mice, CD4^+^ T cell activation was increased in comparison to WT control, and dysregulation of CD8^+^ T cell activation that was dependent upon material treatment (CD4 ECMtx 7.99% active ± 1.22 versus 15.78% ± 2.52, *P* < 0.0001, **Figure 5c,d**). These material-dependent effects followed in peripheral blood at 21 days post-injury, where both CD4^+^ and CD8^+^ T cells had increased activation compared to a WT control (CD4 ECMtx 27.1% ± 15.45 versus 68.02 ± 16.37, *P* = 0.0013, **Figure 5e,f**). Furthermore, at this timepoint there was a lower expression of HELIOS in peripheral blood CD8^+^ T cells (1.83x WT vs *Batf3^-/-^*, *P* < 0.0001) **Figure 5g**). Macrophage polarization was also greatly disrupted at both 7- and 21-days post-injury within the muscle microenvironment, where there was a strong and significant loss of F4/80 expression in all treatment groups (ECMtx 3.05x WT v *Batf3^-/-^*, *P* < 0.0001), along with a modest loss of CD301b in ECMtx, and a prominent loss of CD206 expression (4.02x WT v *Batf3^-/-^*, *P* < 0.0001, **Figure 5h**).

When comparing muscle histopathology at 21 days post-injury, there was a prevalence of hemosiderin in the *Batf3^-/-^*mice suggesting persistent inflammation and blood vessel disruption. Furthermore, muscle fiber shape and morphology were disrupted at the injury interface along with an increase in adipogenesis compared to a wild-type control. In *Batf3^-/-^*ECMtx mice there were a number of stochastically positioned necrotic muscle fibers and potential microcalcifications that were surrounded by cellular infiltrate and giant cells more distal from the injury interface showing a spread of trauma beyond the initial insult (**Figure 5i**).

### Upregulation of XCL-1, E-Cadherin and TGFβ signaling in the injury microenvironment as potential mediators of CD103^+^XCR1^+^ cell recruitment

At 7 days post-injury, an immune infiltration could be detected at the injury interface and around implanted materials. CD103^+^ cells were readily apparent including within clustered areas that could be granulomas or tertiary lymphoid structures (**Figure 6a**). We observed a significant up- regulation of E-Cadherin at the injury when compared to uninjured muscle (fold change: ECMtx = 3.373, PEtx = 3.45, *P* < 0.01), whereas distal uninjured tissue within the injured quad was not significantly different (**Figure 6b,c**). XCL1 protein was higher at 7 days post-injury than 3- and 21- days post-injury in the plasma of injured mice (90.80 pg/ml ± 14.58 vs 63.75 ± 9.93 and 61.04 ± 7.35, *P* < 0.01), and also elevated in local muscle lysate in ECMtx versus PEtx & Control when quantified by ELISA (38.76 ± 9.42 pg/mg protein vs 18.18 ± 2.64 and 25.24 ± 7.11, *P* < 0.05; **Figure 6d,e**). *Rag1^-/-^* mice did not have altered XCL-1 levels in the plasma suggesting the major source is from a RAG-independent cell type (**Supplemental Figure 21**).

**FIGURE 6.**
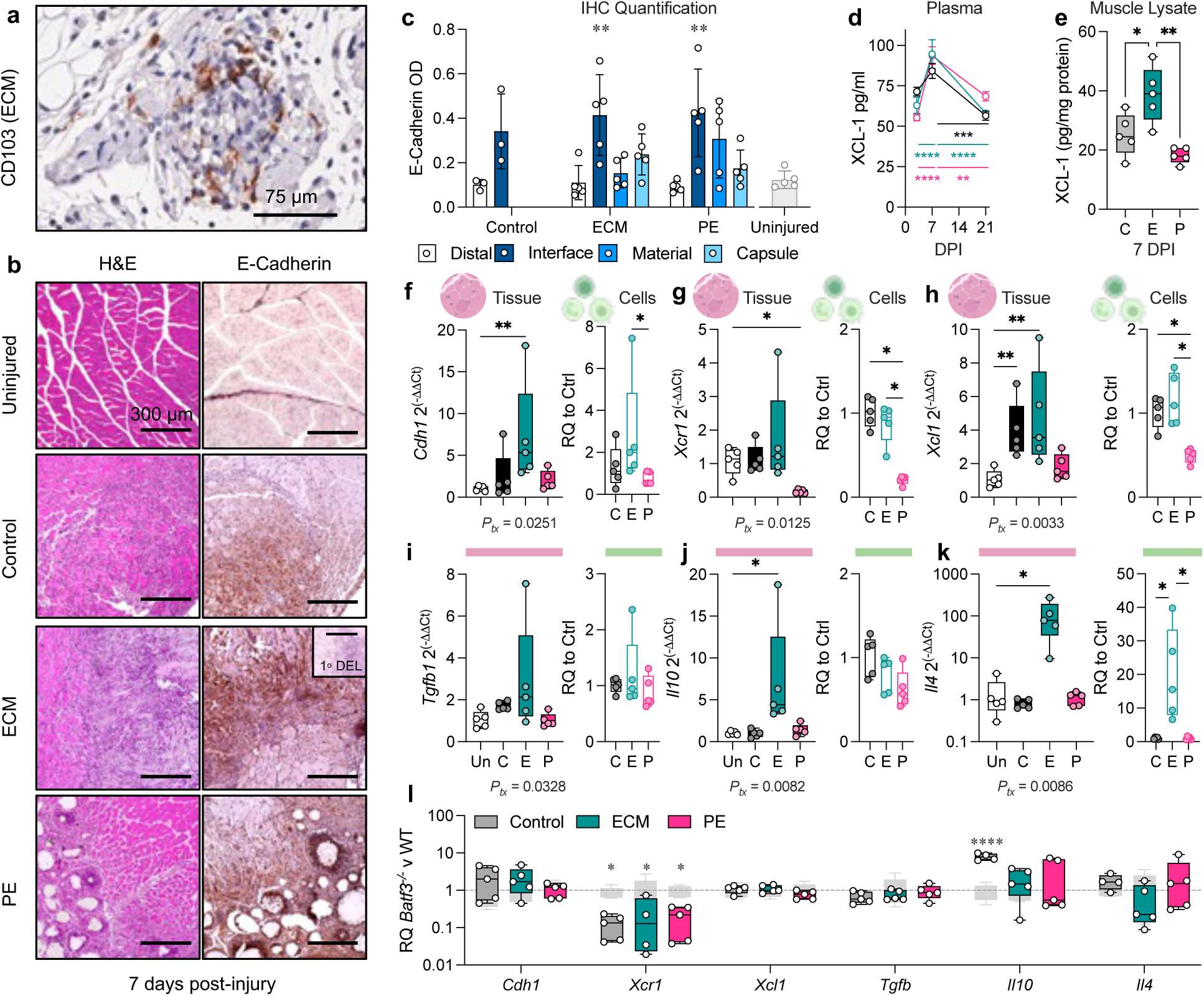
Up-regulation of XCL-1 and E-Cadherin in muscle is associated with tissue damage and material implantation. Gene expression and immunohistochemistry of muscle tissue at 7 days post-injury **(a)** Representative images of hematoxylin and eosin (H&E) and E-Cadherin (DAB) staining at the injury interface at 7 days post-injury. Scale bars = 300μm **(b)** CD103 cells found at the material and injury interface, scale bar = 75μm **(c)** Quantification of E-Cadherin expression via DAB quantification at the distal/uninjured muscle tissue, the injury interface, the material itself, and the external overlaying capsule. N/A = not applicable for control injury or uninjured sample. **(d)** Plasma XCL-1 protein concentrations **(e)** XCL-1 protein concentrations in tissue lysate as w/w over total protein concentration. **(f - k)** mRNA relative quantification as a fold change (2^-ΔΔCt^) over uninjured control for tissue or fold change over control untreated injury for cells isolated from injury, at 7 days post-injury. Kruskall-Wallis with FDR correction for multiple comparisons, * = *P* < 0.05, ** = *P* < 0.01, *Ptx* = the overall significance of injury and treatment on gene expression. **(f)** *Cdh1*, **(g)** *Xcr1*, **(h)** *Xcl1* **(i)** *Tgfb1*, **(j)** *Il10*, **(k)** *Il4*. **(l)** Fold change of *Batf3^-/-^* versus WT gene expression within each treatment condition at 7 days post-injury, grey bars = WT. Data are means ± SEM or range (box plots), *n* = 3 – 5. White = uninjured, Black/grey = Control untreated injury, Teal = ECMtx, Pink = PEtx.

Injury followed by ECM treatment induced an upregulation of *Cdh1* gene expression compared to an uninjured control (*P* = 0.0251; **Figure 6f**). In comparison to an uninjured control, there was significant down-regulation of *Xcr1* in PE-treated muscle injury (*P* = 0.0264), and a significant upregulation of *Xcl1* gene expression in control injury and ECMtx groups when compared to an uninjured control (*P* = 0.0083; **Figure 6g,h**). While no individual treatment was significant, there was a significant difference in *Tgfb1* expression upon injury and material implantation across all treatments (*P* = 0.0328), along with a significant upregulation of *Il10* by ECMtx injuries (*P* = 0.0117; **Figure 6i,j**). We observed a robust up-regulation of *Il4* gene expression by ECM treatment of muscle injuries compared to uninjured muscle (*P* = 0.0309; **Figure 6k**), confirming prior work (*28*). Several of these patterns (including the *Cdh1* and *Il4* upregulation in ECMtx and loss of *Xcr1* and *Xcl1* in PEtx) were repeated in cells isolated from the treated muscle injury when compared to the control injury. Compared to a WT control at 7 days post injury, *Batf3^-/-^* mice had significantly lower *Xcr1* gene expression, but no changes in *Tgfb*, *Cdh1*, *Xcl1*, or *Il4*, and may need to be evaluated later in the time course of injury recovery. With the injury, *Batf3^-/-^*mice had a significantly higher level of *Il10* gene expression by 7 days post-injury (*P* < 0.001), possibly as a compensatory mechanism to immune dysregulation seen in the T cells and macrophages after the loss of cross-presenting dendritic cells (**Figure 6e**).

When evaluating the cytokine and chemokine environment via protein arrays, there were significant differences in the immune profile depending upon the presence of an injury and treatment type. Material implantation was associated with an increase in CCL6, which is produced by neutrophils and macrophages that were recruited by these materials. Both PE and ECM treatment induced myeloperoxidase (MPO) upregulation associated with generating reactive oxygen species (ROS). Interestingly, injury up-regulated Endoglin, a part of the TGFβ receptor, compared to an uninjured control (**Supplemental Figure 22**).

## DISCUSSION

Here we describe the induction of cross-presenting capable DCs by trauma and pro-regenerative material treatment. This is accompanied by MHCII-bearing M2/Mreg macrophages and by CD8^+^ iTregs and ST2^+^ regulatory B Cells, as well as CD103^+^XCR1^+^ adaptive immune cells that are induced by trauma. Recruitment of cross-presenting capable dendritic cells and activation of CD103^+^XCR1^+^ CD8 T cells peaks early during the response to injury and is at a maximum by 7 days post-injury. We detected an up-regulation of both XCL-1 and E-Cadherin within the muscle injury site which may serve as a mechanism for the recruitment of these CD103^+^XCR1^+^ cells through an XCL-1 chemotactic gradient and subsequent binding of CD103 to the upregulated E-Cadherin for migration into the injury site. Low levels of XCL-1 detected in PEtx wounds explains the impaired recruitment of these tDCs to a pro-fibrotic environment. Furthermore, as *Rag1^-/-^* mice had lower levels of CD103 expression on dendritic cells, and TGFβ secreted by Tregs is known to induce CD103 upregulation, this presents a possible mechanism of CD103 upregulation in DCs; however, other cells are known to secrete TGFβ and likely act in concert with adaptive immune cells. As *Rag1^-/-^*mice did not have a defect in XCL-1 levels, we can conclude that much of the XCL-1 production is coming from RAG-independent cells, likely NK Cells which are enriched by pro-regenerative scaffold treatment. In addition, NK cells have previously been described to recruit XCR1-expressing cDC1’s to tumors through XCL-1 (*31*).

These data may suggest a balance of antigen cross-presentation during recovery from wounding modulated by engineered material implantation. In the context of materials that promote type-2 and regulatory immune responses such as decellularized ECM scaffolds, this cross-presentation occurs in an environment that is more amendable to peripheral tolerance and antigen-specific wound repair. In the context of pro-inflammatory and pro-fibrotic materials such as PE, which favor a type-1 and type-17 immune response, this could lead to autoimmune activation and formation of auto-reactive T cells and antibodies leading to distal pathologies and systemic immune dysregulation as has been reported in some patients (*32*). This also agrees with previous findings on regulatory CD8^+^ T cells in respiratory infection and laboratory models of autoimmune disease (*20, 21*). In the absence of tDCs in a *Batf3^-/-^* mouse model, increased inflammation and tissue destruction including distal necrosis and calcifications of muscle tissue suggest a continued and spreading pathology after the initial muscle loss. Additionally, increased adipogenesis at the injury site in *Batf3^-/-^* mice suggests a defect in fibro-adipogenic lineage commitment and impaired muscle regeneration.

Presence of CD103^+^XCR1^+^ innate and adaptive immune cells may present a homeostatic regulation of response to injury that is expanded during trauma after reaction with cross-presenting capable dendritic cells (**Supplemental Figure 23**). Previous work has shown that XCR1^+^ T cells can be induced through trogocytosis and communication with cross-presenting capable dendritic cells and they are a potential targets for cancer immunotherapy (*33*). As CD103^+^XCR1^+^ adaptive immune cells were not up-regulated in cross-presentation deficient *Batf3^-/-^* mice, this may suggest a communication between innate and adaptive immune cells that express CD103 and XCR1. In terms of antigen specificity, while the ECM scaffold introduces a protein source for new exogenous antigens, both the control injury and the PE-treated injury are surgically induced sterile trauma that does not introduce non-self-antigen. Thus, these cells are likely reacting to self-antigen, or in an antigen-independent manner. The increase in T cell activation, decrease in HELIOS expression, decrease in ST2^+^ regulatory B Cells, decrease in both F4/80 and CD206 on scaffold-associated macrophages, and expanded tissue damage in *Batf3^-/-^* mice that lack tDCs support our hypothesis that these cells are acting in a regulatory or tolerogenic manner.

To our knowledge, the recruitment and modulations of these and their communication with cross-presenting capable dendritic cells in the context of trauma and biomaterial implantation have not been previously described and constitute a novel mechanism of immune response to wounding and damaged self in traumatic injury, that has conservation of signaling molecules between mice and humans (*34*). While this is likely not the only mechanism of peripheral tolerance at a wound site, it appears to be a major pathway for regulating of inflammation after injury and preventing continued tissue damage after the initial insult. Future work will focus on identifying the source of the stimulation – be it self-antigen, microbiome-associated, or antigen-independent – and the development of therapeutics to modulate these pathways and promote tissue regeneration and medical device integration.

## ACKNOWLEDGEMENTS

The authors would like to thank R Germain, Y Belkaid, M Wolf, and B Warner for helpful conversations. This work was funded by the intramural research program of the National Institutes of Health, National Institute of Biomedical Imaging and Bioengineering, National Institutes of Health (NIH).

## Disclaimer

The contents of this publication are the sole responsibility of the authors and do not necessarily reflect the views, opinions, or policies of the NIH and the Department of Health and Human Services (HHS). Mention of trade names, commercial products, or organizations does not imply endorsement by the U.S. Government.

## AUTHOR CONTRIBUTIONS

RL, TBN, SD, KMA, MB, AJ, AL, MF, MK, PF, and KS performed experiments. RL, TBN, PF, and KS analyzed data. RL and KS wrote the manuscript. RL, TBN, MK, and KS edited the manuscript. KS provided funding and supervision.

## MATERIALS AND METHODS

### Material Preparation

Small intestine was sourced from 5- to 6-month-old American Yorkshire pigs (Wagner Meats). The submucosa layer (SIS) was mechanically isolated by removing muscularis layer and subsequent mechanical scraping of the luminal layer. The resulting SIS was rinsed in distilled water followed by distilled water with 1% anti-biotic and anti-mycotic (penicillin, streptomycin, amphotericin B, Gibco) and frozen at -80°C until decellularization. After thawing, within a biosafety cabinet SIS was cut into 1-inch segments and then incubated in 4 % ethanol (Fisher Scientific) and 0.1 % peracetic acid (Sigma) for 30 minutes with vigorous shaking or on a stir plate. The decellularized ECM was neutralized with successive washes of sterile 1xPBS and distilled water. Liquid was blotted with sterile absorbent pads then the material was transferred to a 50 ml conical tube and frozen at -80°C until lyophilization for 48 hours or until the tissue was dried completely. The dried material was loaded into sterile cryogenic milling containers and milled into a fine powder using a SPEX SamplePrep cryogenic milling device. The resulting powder was hydrated with sterile saline to form a thick paste that was then loaded into a slip-tip 1 ml syringe for application into the wound site.

Polyethylene powder (PE) particle size < 150 μm was purchased from Goodfellow Cambridge Limited and soaked in distilled water before rinsing with 70% ethanol and UV sterilized in ethanol for 30 minutes. PE was stored in 70% ethanol until use. PE particles suspended in ethanol were transferred to an Eppendorf tube and dried overnight in a biosafety cabinet. Due to the hydrophobicity of PE, samples cannot be loaded into a syringe and are applied directly to the wound as a powder.

### Volumetric Muscle Loss Surgery

Mice received bilateral volumetric muscle loss trauma as per the previously described method (*26*). Briefly, the lower limbs of 5 to 8-week-old female C57BL/6 WT, *Rag1^-/-^*, or *Batf3^-/-^*mice (Jackson Laboratory stocks: 000664, 002216 (*35*), 013755 (*36*)) were shaved with an electric razor and cleared of excess hair by depilatory cream, one day before surgery. The following day, mice were anesthetized in anesthesia chamber under 4.0% isoflurane in oxygen at a 200 cc/min flow rate and received subcutaneous injection of slow-release buprenorphine for pain management (0.5 mg/ml). The mice were then maintained at 2.0% isoflurane for the duration of the procedure which concluded in under 10 minutes. The surgical site was sterilized with three rounds of betadine followed by 70% isopropanol prior to making a 1 cm incision in the skin and through the fascia above the quadriceps muscles. Using surgical scissors, a 3–4 mm defect was created in the mid-belly section of the quadriceps muscle by removing 1/3 of the quadriceps muscle. After muscle removal, resulting tissue gap was filled with uniform amount (50 μl) of either polyethylene particulate, porcine-derived ECM scaffold, or a saline vehicle control. The wound was then subsequently closed with 3-4 wound clips (7 mm, Roboz) and the procedure was repeated on the contralateral leg. After the surgery mice were placed under a heat lamp for 2-3 minutes to let them recover from anesthesia and surgical tools were sterilized in a glass bead sterilizer. Mice were then placed back in the cage and monitored closely until ambulatory and grooming. They were then maintained on regular diet with enrichment until the end of the study. The protocol was approved by the NIH Clinical Center Animal Care and Use Committee under animal protocol number NIBIB 20-01.

### Flow cytometry

After 3-, 7-, 21- and 42-days post-injury, mice were ethically euthanized followed by dissection of injured muscle along with implanted scaffolds. The dissected muscle was then finely diced and for the myeloid panel, digested with digestive media (0.5 mg/mL Liberase TM (Sigma) and 0.2 mg/ml DNase I (Roche) in HEPES supplemented media) on a shaker at 100 rpm for 45 minutes and 37°C. The digested suspension was then filtered through a 70 µm cell strainer and washed with 1x PBS and centrifuged at 350g for 5 minutes at room temperature. The cell pellets were then soaked for 10 minutes in 5mM EDTA in 1xPBS solution to reduce cell clumping. After, 10 minutes of incubation, cells were then again washed with 1 x PBS followed by centrifugation at 350 g for 5 minutes at 4°C. For the lymphoid panel, diced muscle was homogenized through 70 μm cell strainers (lymph node samples) or in a mechanical homogenizer (Fisher) briefly in 5mM EDTA in 1xPBS prior to centrifugation and isolation of a cell suspension, this was due to sensitivity of lymphoid markers (CD4 & CD8) to the digestive enzymes used in the myeloid panel. For blood samples, blood was collected in a K2EDTA microtainer through cheek or retroorbital bleed, then briefly vortexed, centrifuged to remove plasma, then resuspended in RBC lysis buffer (Qiagen) for 10 minutes on ice before centrifugation and washing the cell pellet with 1xPBS.

The cell pellets were then re-suspended in 200 µL viability dye solution (1:1000 dilution of Live/Dead Blue (Thermo Fisher) in PBS) for 20 minutes on ice followed by washing with wash buffer (1% BSA and 2mM EDTA in 1x PBS). The cells were stained with either myeloid panel or lymphoid panel antibodies (**Supplementary Table 1,2; Supplemental Figures 24 - 26**) followed by incubation at 4 °C for 30 min. After incubation the cells were washed three times with wash buffer and analyzed on a 5 Laser Cytek Aurora flow cytometer.

### Intracellular staining

After surface staining, cells were fixed and permeabilized by using True-Nuclear™ Transcription Factor kit (BioLegend) for intracellular staining of FoxP3 and HELIOS antibodies (**Supplementary Table 2**) as per manufactured guidelines. Briefly, after last washing of surface staining, cells were resuspended in True-Nuclear™ 1x Fix Concentrate and incubated for 45 minutes at 4 °C. After incubation the cells were centrifuged at 400xg for 10 minutes at 4°C and resuspend in True- Nuclear™ 1x Perm Buffer. The cells were then again washed for one additional time with 1x Perm Buffer and were then treated with HELIOS and FoxP3 antibody cocktail (1:100 dilution of antibodies in True-Nuclear™ 1x Perm Buffer) followed by incubation at 4°C for 45 minutes. After incubation cells were washed twice with True-Nuclear™ 1x Perm Buffer. After final wash with Perm Buffer, the cells were re-suspended in wash buffer (1% BSA and 2mM EDTA in 1x PBS) and were analyzed on a 5 Laser Cytek Aurora flow cytometer.

### Histopathology

Samples were fixed in 10% neutral buffered formalin for 48 – 72 hours prior to transfer to 70% ethanol. Samples were then dehydrated in graded ethanol steps through 70%, 80%, 95%, and 100% ethanol prior to clearing in xylene and embedding in paraffin wax using an automated tissue processor (Leica Biosystems). Quadriceps muscle groups were then cut in a transverse fashion to expose the center of the injury, which was then mounted face down in the paraffin mold. Five (5) to 7 μm sections were then placed onto charged glass slides and baked overnight at 56°C to dry. After rehydration, samples were stained with hematoxylin and eosin (H&E) through a 5- minute incubation in Harris Hematoxylin (Sigma Aldrich), followed by a wash in tap water and three dips in acid ethanol to destain, prior to rinsing in ethanol and dipping in Eosin Y (Sigma Aldrich). Samples were then washed in 95% ethanol, dehydrated, coverslipped and mounted with Permount. Slides were imaged on an EVOS microscope (Thermo).

### Immunohistochemistry

Samples were rehydrated, then incubated for 20 minutes in citrate antigen retrieval buffer prior to slowly cooling for 20 minutes on the benchtop. Endogenous peroxidases were quenched through a 5-minute incubation in 0.3% hydrogen peroxide in 1xPBS. Samples were stained using the VECTASTAIN® Elite ABC-HRP Kit (Rabbit, Vector Laboratories) as per manufacturer’s instructions. Briefly, after washing in 1xPBS, samples were blocked in 2.5% normal goat serum for 1 hour. Samples were incubated in primary antibody diluted in blocking buffer for 1 hour. Rabbit monoclonal anti-CD103 (AbCam) and anti-E-Cadherin (AbCam) were diluted at a 1:100 dilution and 1:5000 dilution, respectively. Slides were washed 3 times in 1xPBS prior to incubation with biotinylated secondary antibody for 30 minutes. Slides were washed 3 times in 1xPBS then incubated for 30 minutes in the VECTASTAIN Elite ABC Reagent. Samples were washed 3 times in 1xPBS, and then incubated in ImmPACT® DAB EqV Peroxidase (HRP) Substrate (Vector Laboratories) for 1 minute and 30 seconds (CD103) or 60 seconds (E-Cadherin). Slides were washed in tap water, then counterstained for 5 minutes in Harris Hematoxylin (Sigma), prior to rinsing in tap water, destaining in acid ethanol, dehydration and mounting in Permount.

### RNA isolation and RT-PCR

Quadriceps muscle group was dissected from mice at 7 days post-injury and homogenized in 2 ml 1xPBS with a mechanical homogenizer at 5000 rpm for 30 seconds. Five hundred (500) microliters of the resulting homogenate was transferred to an Eppendorf tube containing 500μl of TRI Reagent Solution (Sigma Aldrich). Samples were vortexed and then stored at -80C until RNA isolation. After thawing, 200μl of chloroform (Sigma Aldrich) was added to each sample and vortexed before being allowed to separate for 5 minutes at room temperature, followed by centrifugation for 15 minutes at 8000 xg and 4°C. Aqueous phase was combined with an equal volume of 70% ethanol and vortexed. The resulting sample was passed through an RNeasy Mini Prep spin column (Qiagen), and then washed as per manufacturer’s instructions with 1 wash of RW1 Buffer, and 2 washes of RPE Buffer. The membrane was dried, and RNA was eluted into 30ul of RNAse-free water. NanoDrop was used to determine RNA concentrations and quality control was performed to move forward with samples with an A260/A280 > 2. Samples were diluted to 150 ng/μl concentration, and 11 μl of each were added to a SuperScript Reverse Transcriptase IV reaction following manufacturer’s instructions with Random Hexamers as primers (ThermoFisher Scientfic). Two (2) μl of the resulting cDNA was added alongside 10 μl of TaqMan Fast Advanced Master Mix, 7 μl of nuclease-free water, and 1 μl of FAM-MGB primer/probe: *Gusb*, *Cdh1, Xcr1, Xcl1, Il4, Il10, Tgfb1* (See **Supplemental Table 3** for details). Samples were then run on an Applied Biosystems QuantStudio 3 RT-PCR Machine (ThermoFisher).

### Protein Isolation from mouse tissue and cytokine analysis

Muscle and lymph node samples were flash frozen in liquid nitrogen or an ethanol-dry ice slurry immediately after dissection and stored until processing. Frozen muscle samples were added to 2 ml ice cold 1xPBS with protease inhibitors (ThermoFisher Scientific) and diced with a pair of scissors. Samples were homogenized for 45 seconds using a mechanical homogenizer at 5000 – 6000 rpm while on ice. Subsequently, 2.5 ml more ice cold 1xPBS with protease inhibitors were added along with 50μl of 10% Triton-X100 then mixed vigorously and left on ice for 5 minutes prior to aliquoting and snap freezing in liquid nitrogen and stored until use. Day of use, samples were thawed and centrifuged at 10,000xg for 10 minutes to pellet debris. Protein concentration was determined via the Pierce BCA Protein Assay Kit (Thermo Scientific) at a 1:1 dilution with lysis buffer. Two hundred (200) micrograms of protein were loaded onto a Proteome Profiler^TM^ Array, Mouse XL Cytokine Array Kit and assayed per manufacturer’s instructions. Blots were imaged on a BioRad ChemiDoc with a 30 second exposure.

### Enzyme-linked immunosorbent assay

The XCL-1 measurement in mouse blood plasma samples was performed by using Mouse XCL- 1 SimpleStep ELISA kit (Abcam). The assay was performed as per manufacturer guidelines. Briefly, 50µL of 1:1 diluted mouse blood plasma sample: blocking buffer was added to appropriate wells of precoated 96 well plate. The samples were then treated with 50µL of antibody cocktail followed by incubation for 1 hour at room temperature. After incubation, the mixture in each well was aspirated and wells were washed three times with wash buffer. After final wash, 100µL of TMB development solution was added to each well and plate was incubated for 10 minutes. After incubation, 100µL of stop solution was added to each well followed by reading OD at 450nm. Protein lysate was loaded to the ELISA plate 1:1 with dilution buffer and then normalized to total protein concentration determined in the BCA as previously mentioned and proceeded through manufacturer’s protocols.

### Statistics and Data Analysis

Flow cytometry data were unmixed using stated single spectra controls (Supplemental Tables 1,2) using SpectroFlo Software (Cytek Biosciences). Resulting unmixed data were exported to .fcs prior to analysis on FlowJo (Supplemental Figures 2,16). Dimensionality reduction algorithms were run through FlowJo plugins. T-stochastic neighbor embedding (t-SNE) was run at the following parameters: learning configuration – opt-SNE, iterations – 2000, perplexity – 30, KNN algorithm – exact (vantage point tree), gradient algorithm – Barnes-Hut (*37*). Uniform manifold projection was run at the following parameters: Euclidean, nearest neighbors – 15, minimum distance – 0.5, Number of components – 2 (*38*). FlowSOM 3.0.18 was run at the following parameters: Number of meta clusters – 30 (*39*). Clustering was run on singlet live immune cells using all parameters excluding LIVE/DEAD Blue, for myeloid panel, and CD11b-CD11c- singlet live immune cells excluding LIVE/DEAD Blue and CD11b/c, for lymphoid panel. Resulting data from manual gating and dimensionality reduction algorithms were analyzed and displayed using GraphPad Prism v9 and R 4.1.2.

Immunohistochemistry of E-Cadherin was quantified through auto-white balance in Fiji (ImageJ) followed by Color Deconvolution to isolate the DAB channel. Areas of interest were manually outlined then measured, and transformed into optical density (OD) readings by taking the log(max intensity/Mean intensity). Each replicate represents quantification of a section from a different animal. Resulting OD values were plotted in GraphPad Prism v9 for data display and analysis.

Chemilumiescent proteome profiler blots were quantified by pixel intensity via MatLab (version R2022a) using the Protein Array Tool version 2.0.0.1 and normalized to background prior to being displayed as a fold change over uninjured control muscle tissue.

RT-PCR was calculated as a fold change over uninjured (tissue) or control injury (cells from tissue) through the 2^-ΔΔCt^ method with *Gusb* as the housekeeping gene. In the case where a treatment did not yield detectable signal above threshold by 40 cycles of PCR, 40 was inserted as the minimum cycle estimate and fold change was calculated as the minimum potential difference in gene expression. For samples in which there was evidence of primer dimer or self-replicating hairpin contamination determined by an early amplification that crossed threshold and plateaued quickly at a low intensity (<10 cycles), these samples were excluded from analysis.

Statistical tests used are stated in figure captions. If data were normally distributed, ANOVA with Tukey post-hoc correction for multiple comparisons was used. If data were not normally distributed, Kruskall-Wallis with FDR post-hoc correction for multiple comparisons was used.

## SUPPLEMENTAL FIGURES AND LEGENDS

**Supplemental Figure 1.**
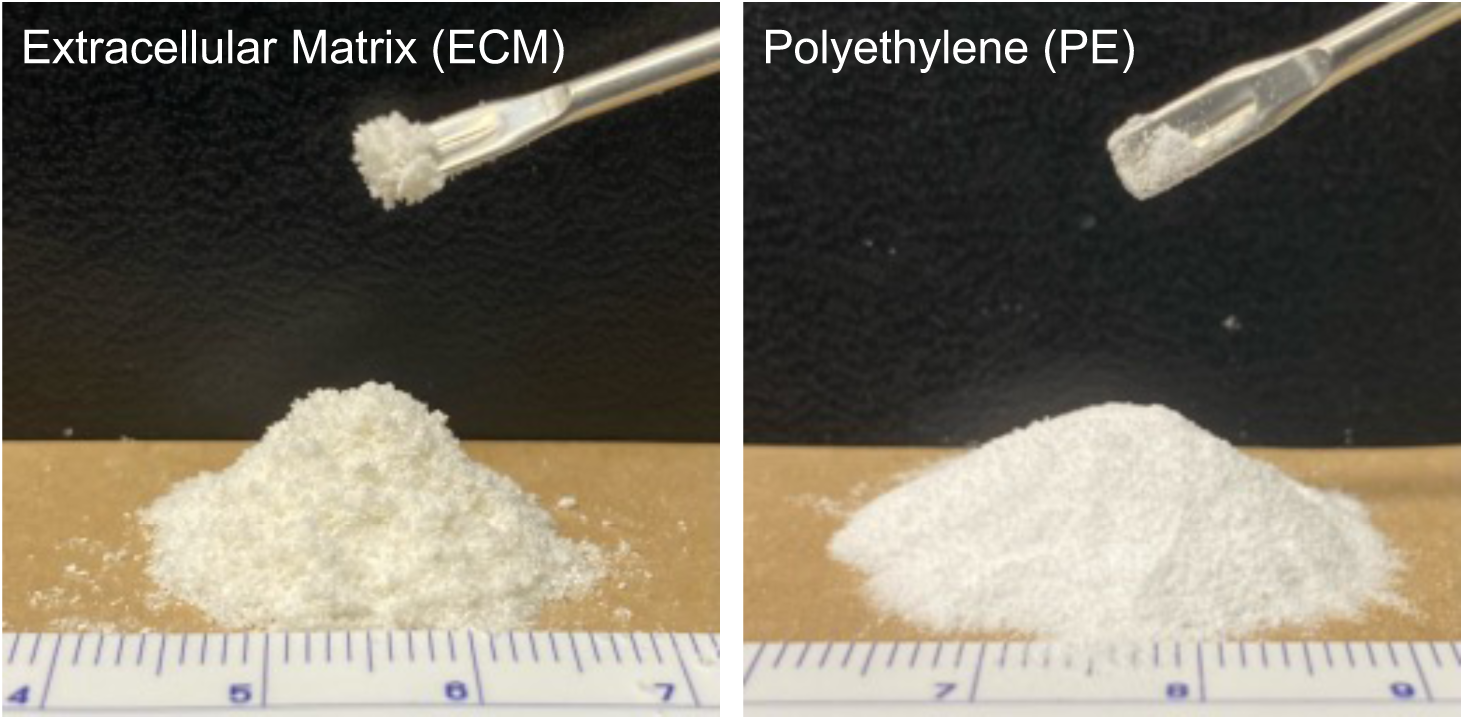
Materials used. Macroscopic image of ECM powder and PE powders.

**Supplemental Figure 2.**
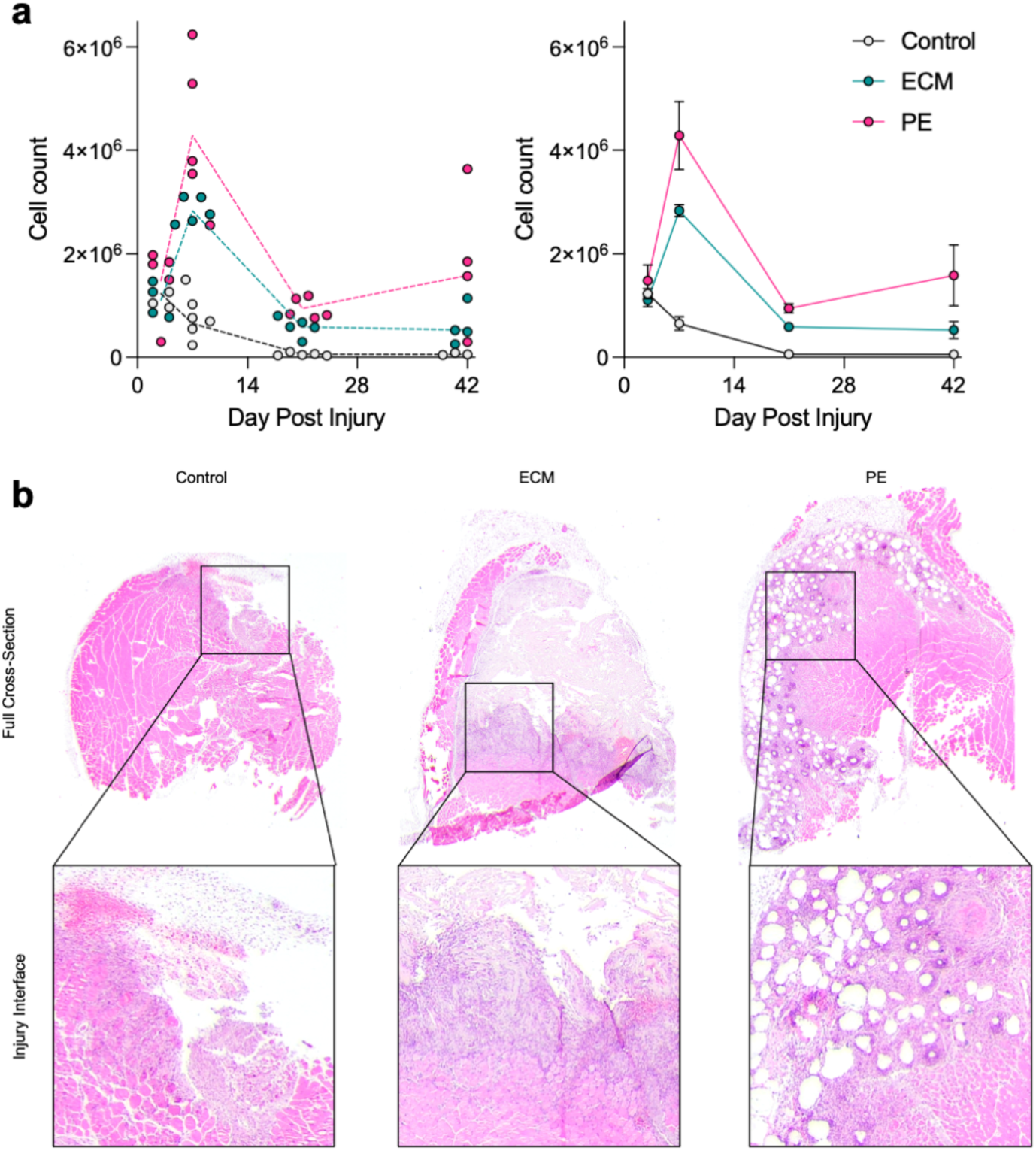
Immune cell infiltration into muscle injury. (a) Cells were counted on a hemocytometer prior to flow staining, live immune cell counts are displayed. Left individual values per mouse, right = mean ± SEM. (b) Hematoxylin and eosin staining of muscle injury at 7 days post-injury.

**Supplemental Figure 3.**
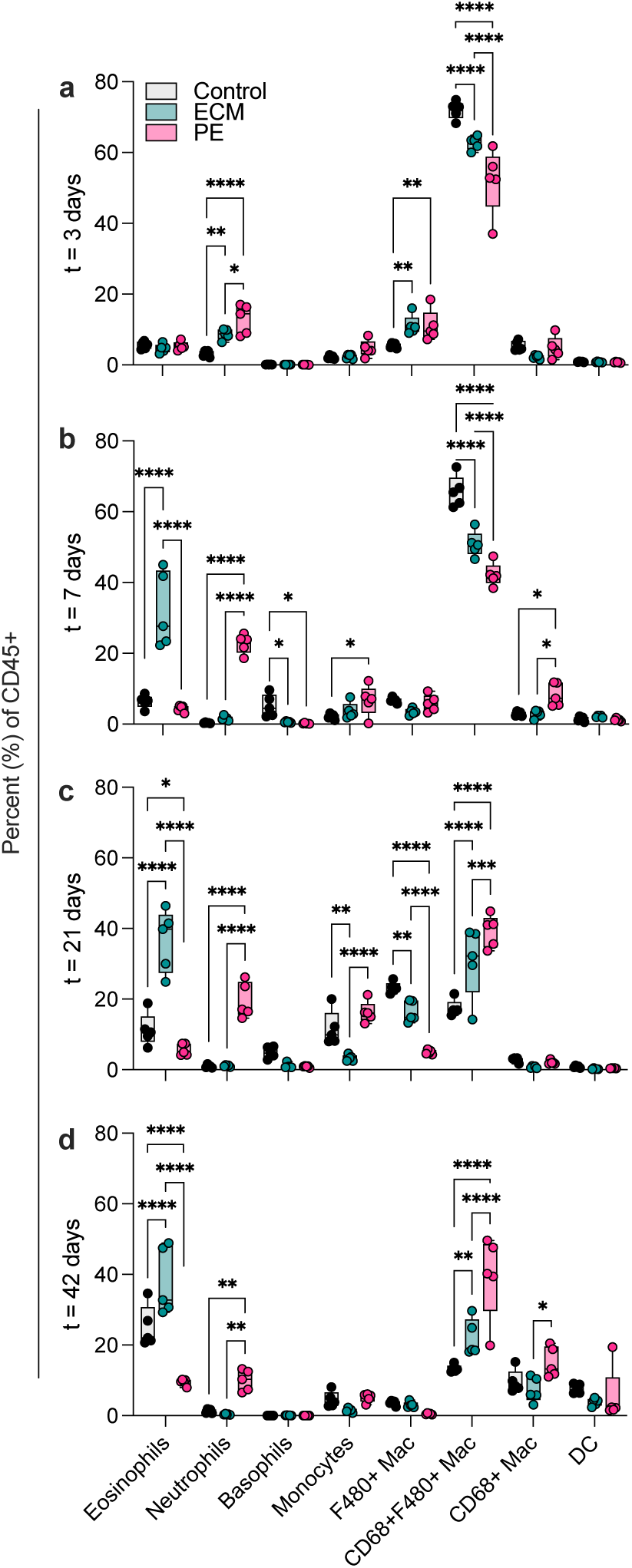
Innate immune cell phenotyping in trauma microenvironment. (a) 3 days post-injury (b) 7 days post injury (c) 21 days post injury (d) 42 days post-injury. Control = black, ECM = teal, PE = pink. ANOVA with Tukey post-hoc correction for multiple comparisons, * = *P*<0.05, ** = *P*<0.01, ***= *P*<0.001, ****= *P*<0.0001. *n* = 5.

**Supplemental Figure 4.**
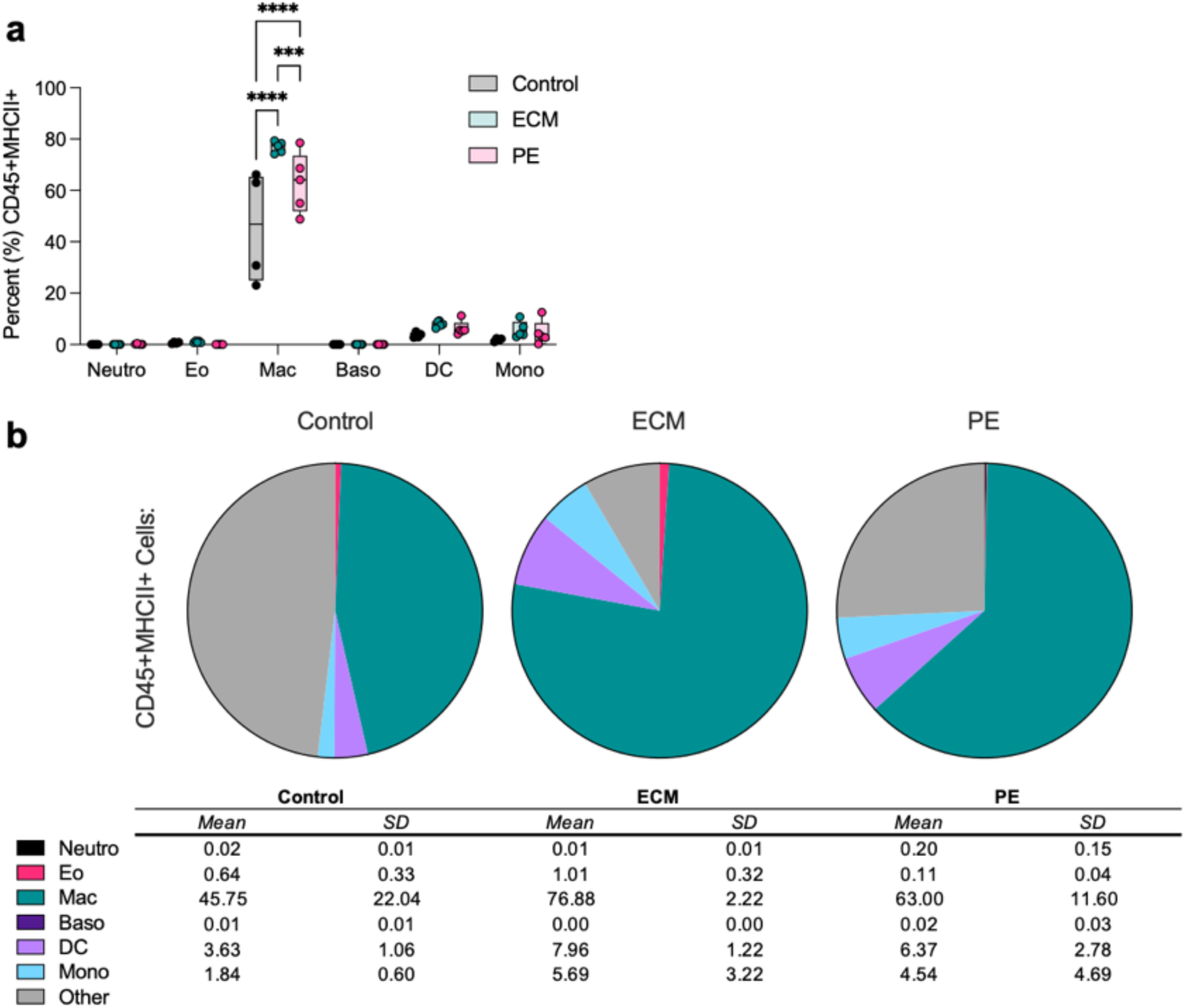
Phenotyping of MHCII^+^ antigen presenting cells in the wound microenvironment at 7 days post injury. (a) Percent of MHCII^+^ cells that are identified as listed immune cell types (b) Proportion of each treatment group that is represented by different cell types. SD = standard deviation. Black = neutrophils, pink = eosinophils, teal = macrophages, purple = basophils, lavender = dendritic cells, blue = monocytes, grey = other immune cells. ANOVA with Tukey Post-hoc correction for multiple comparisons, * = *P*<0.05, ** = *P*<0.01, *** = *P*<0.001, **** = *P*<0.0001.

**Supplemental Figure 5.**
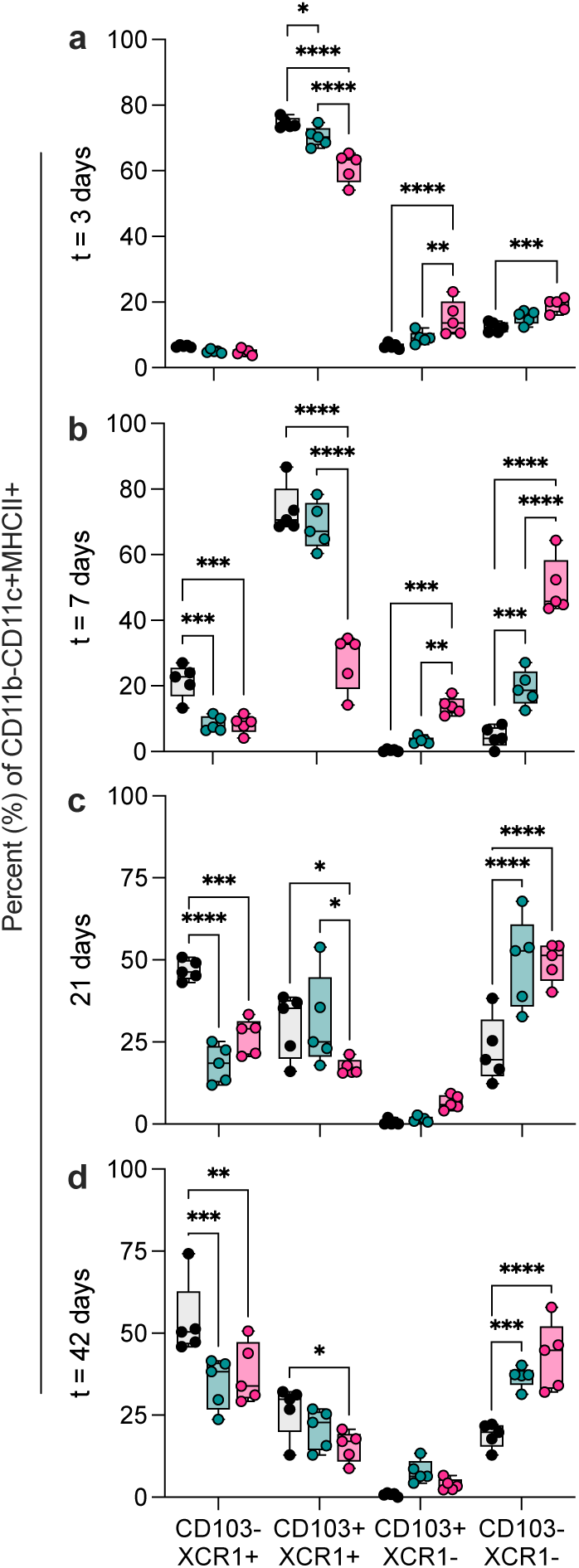
Dendritic cell subsets at (a) 3 days post-injury (b) 7 days post-injury, (c) 21 days post injury, (d) 42 days post-injury. ANOVA with Tukey Post-hoc correction for multiple comparisons,* = *P*<0.05, ** = *P*<0.01, *** = *P*<0.001, **** = *P*<0.0001. Data are means ± SEM, *n* = 5 representative of at least two independent experiments.

**Supplemental Figure 6.**
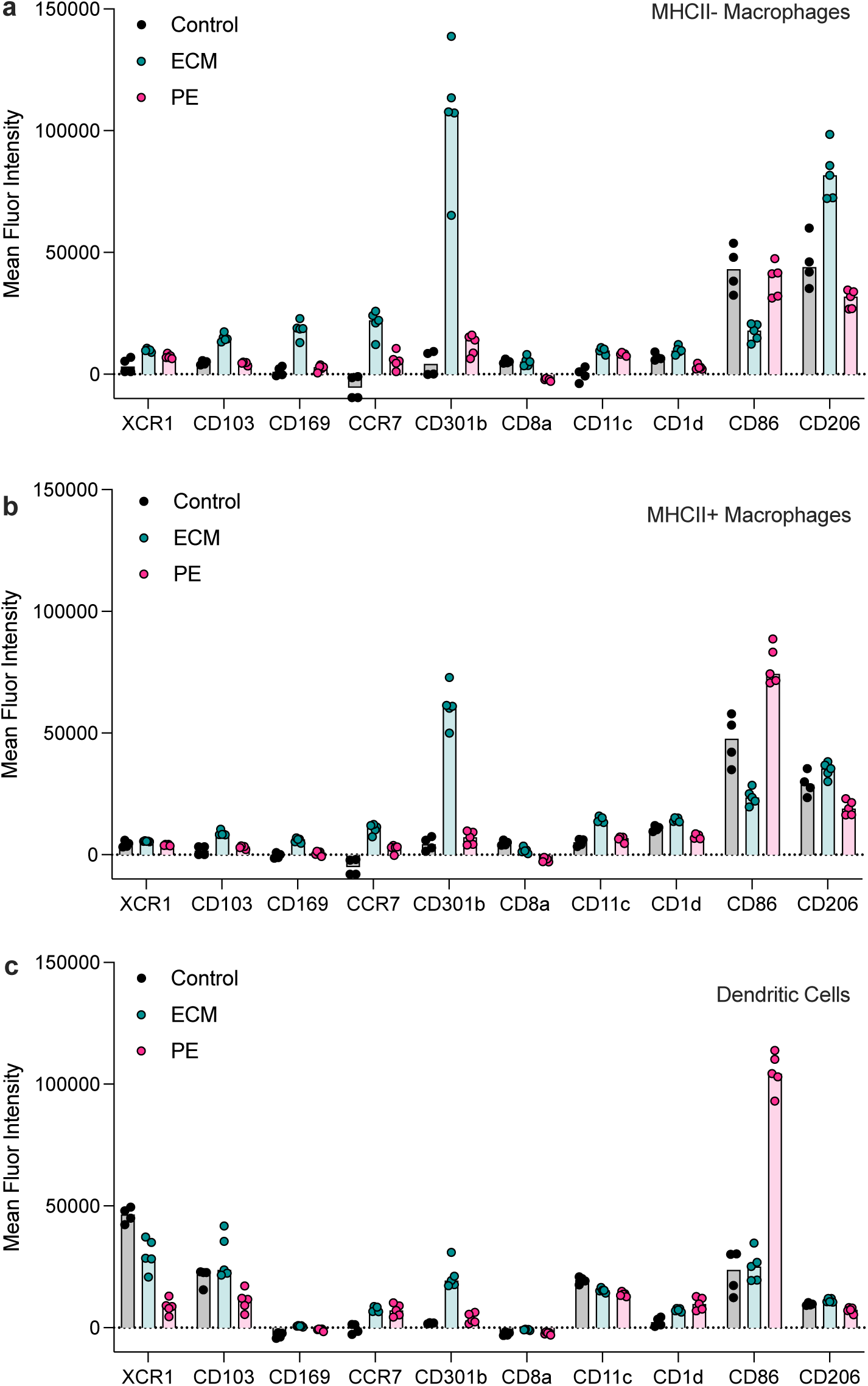
Phenotyping markers of macrophages and dendritic cells. (a) MHCII^+^ macrophages (F4/80^+^CD68^+^MHCII^+^). (b) MHCII^-^ Macrophages (F4/80^+^CD68^+^MHCII^-^). (c) Dendritic cells (CD11c^+^MHCII^+^). Black/grey = control injury; teal = ECM-treated injury; pink = PE-treated injury. *n* = 5 data are shown as individual points with bar at mean.

**Supplemental Figure 7.**
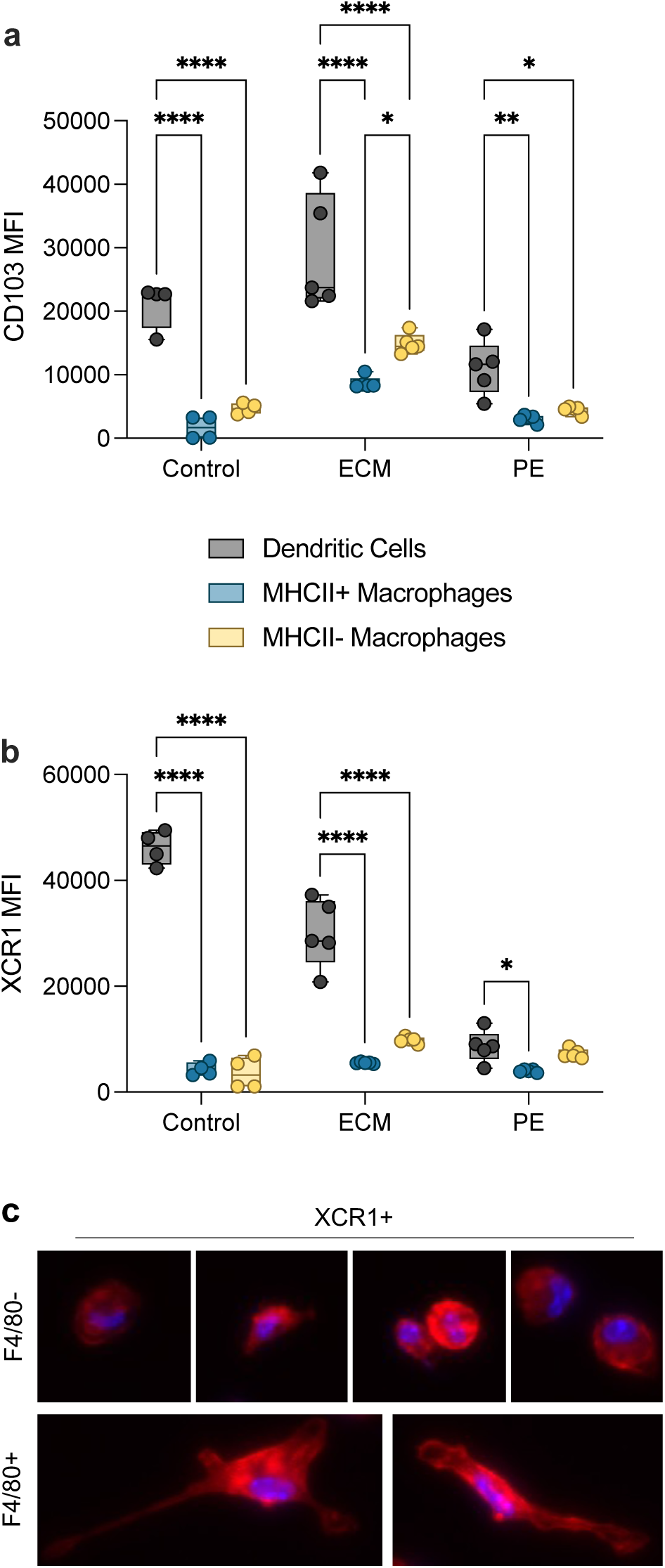
CD103 and XCR1 expression on myeloid cells. (a) CD103 mean fluorescence intensity, and (b) XCR1 mean fluorescence intensity on dendritic cells (black), MHCII+ macrophages (blue), and MHCII- macrophages (yellow). (c) F4/80^-^ and F4/80^+^ fractions of XCR1^+^ cells purified (via MACS column) from an ECM-treated VML at 7 days post-injury. Red = actin/phalloidin, Blue = DAPI. *N* = 4 – 5, ANOVA with Tukey post-hoc correction for multiple comparisons, * = *P* < 0.05, ** = *P* < 0.01, **** = *P* < 0.0001.

**Supplemental Figure 8.**
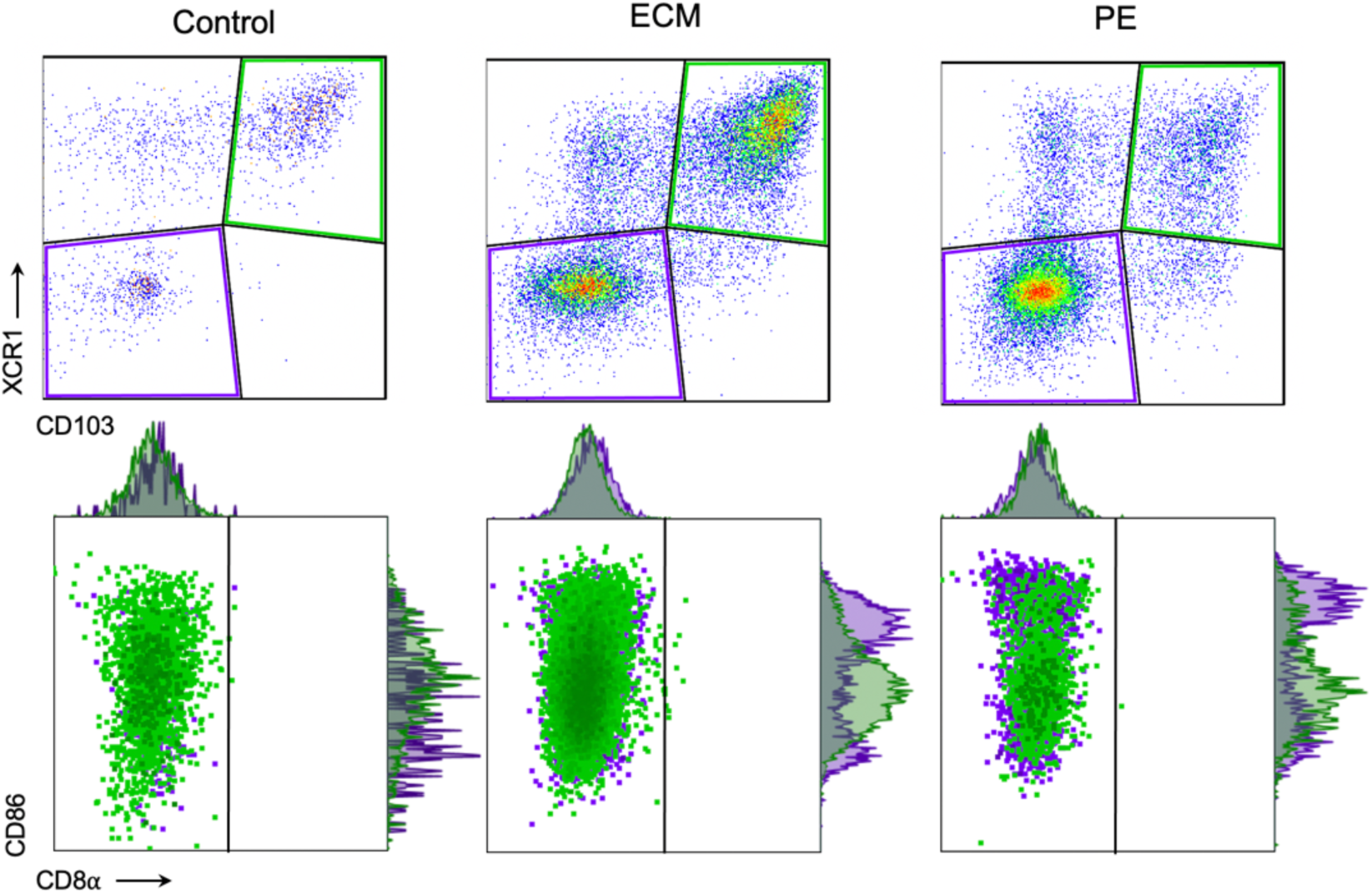
CD86 expression on tDCs. Representative FACS plots of CD11c^+^MHCII^+^ dendritic cells in three treatment groups. Green = XCR1^+^CD103^+^ tDCs; purple = XCR1^-^CD103^-^ dendritic cells.

**Supplemental Figure 9.**
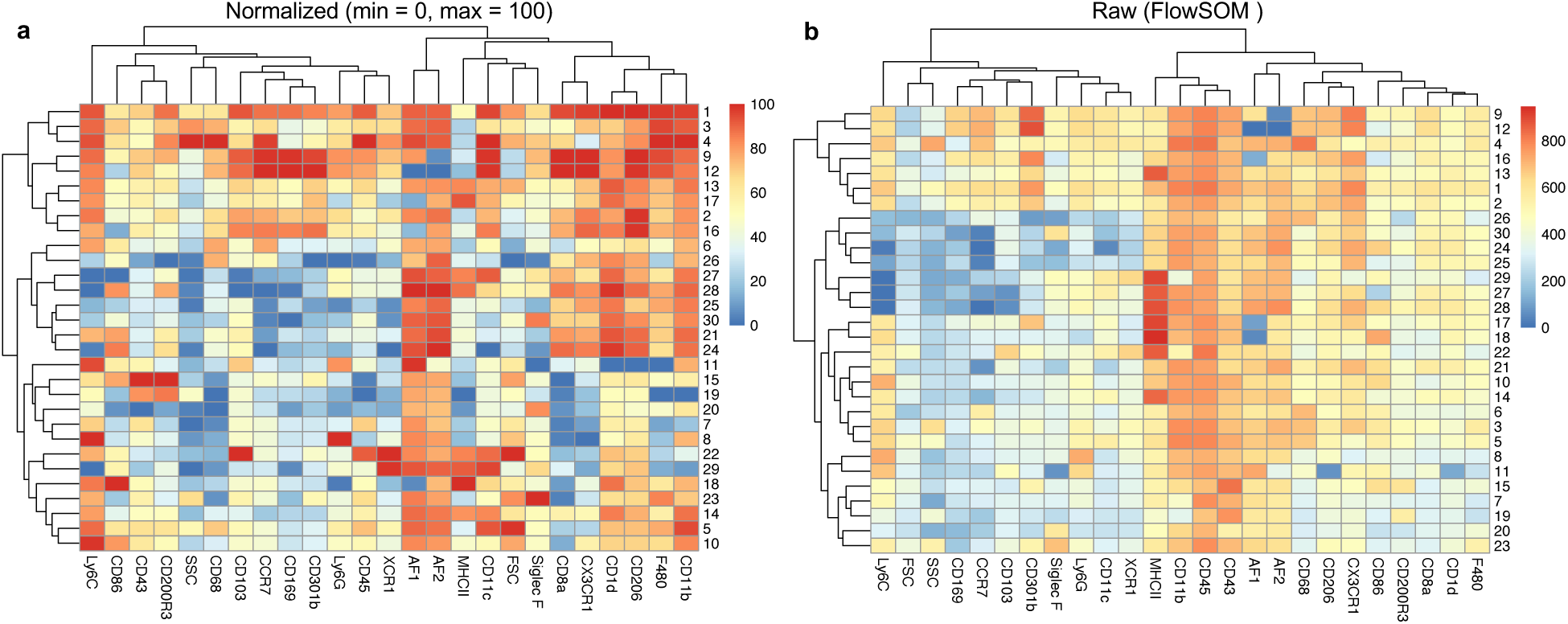
Phenotyping and hierarchical clustering of FlowSOM-derived cell populations. (a) Normalized FlowSOM data min = 0, max = 100. (b) Non-normalized FlowSOM data (scaled).

**Supplemental Figure 10.**
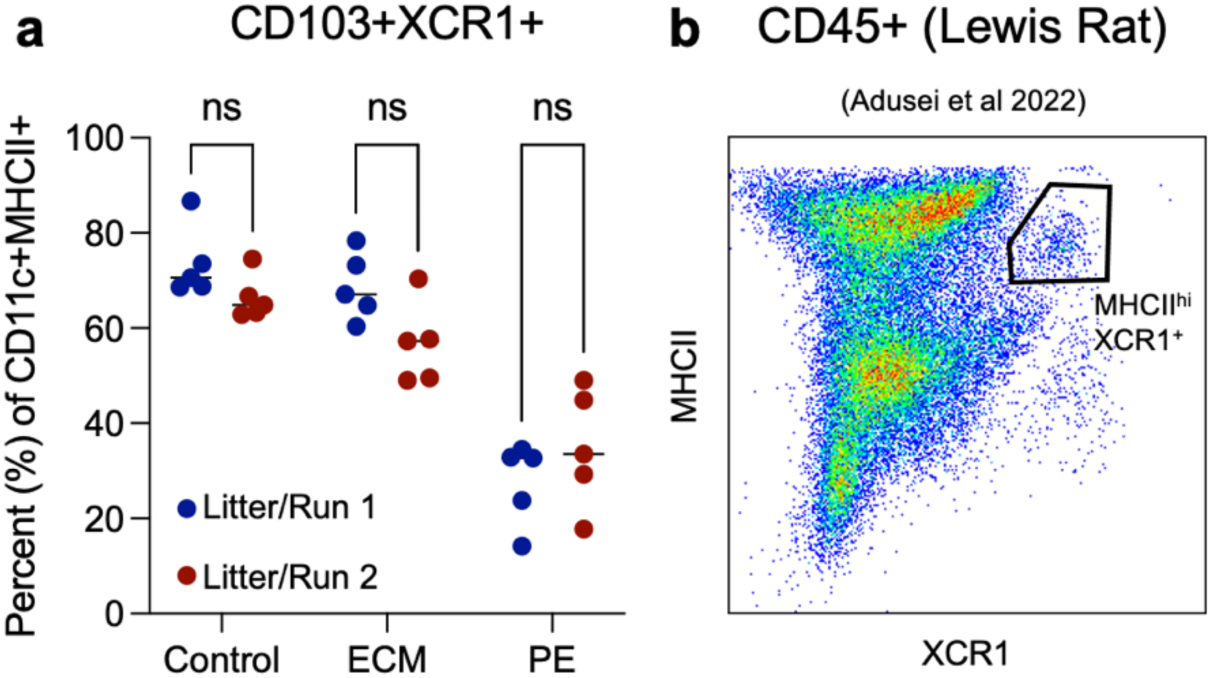
Repeatability of findings across litters and species. XCR1^+^CD103^+^ cells are present in multiple runs with mice from (**a**) different litters and (**b**) species. Student’s T-test with Tukey post-hoc correction, ns = not significant. Rat data are extracted from Adusei et al, *Cells, Tissues, and Organs*, 2022.

**Supplemental Figure 11.**
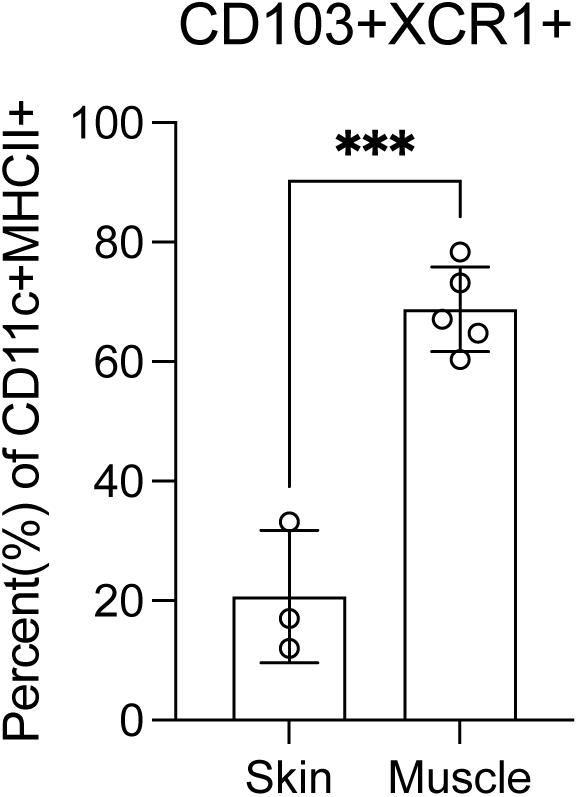
CD103^+^XCR1^+^ Dendritic cells in the skin overlying a muscle injury. At 7 days post-injury in an ECM-treated mouse, as a proportion of CD11c^+^MHCII^+^ dendritic cells. Data are means ± standard deviation, *n* = 3 - 5. Student’s t-test, *P* = 0.0003.

**Supplemental Figure 12.**
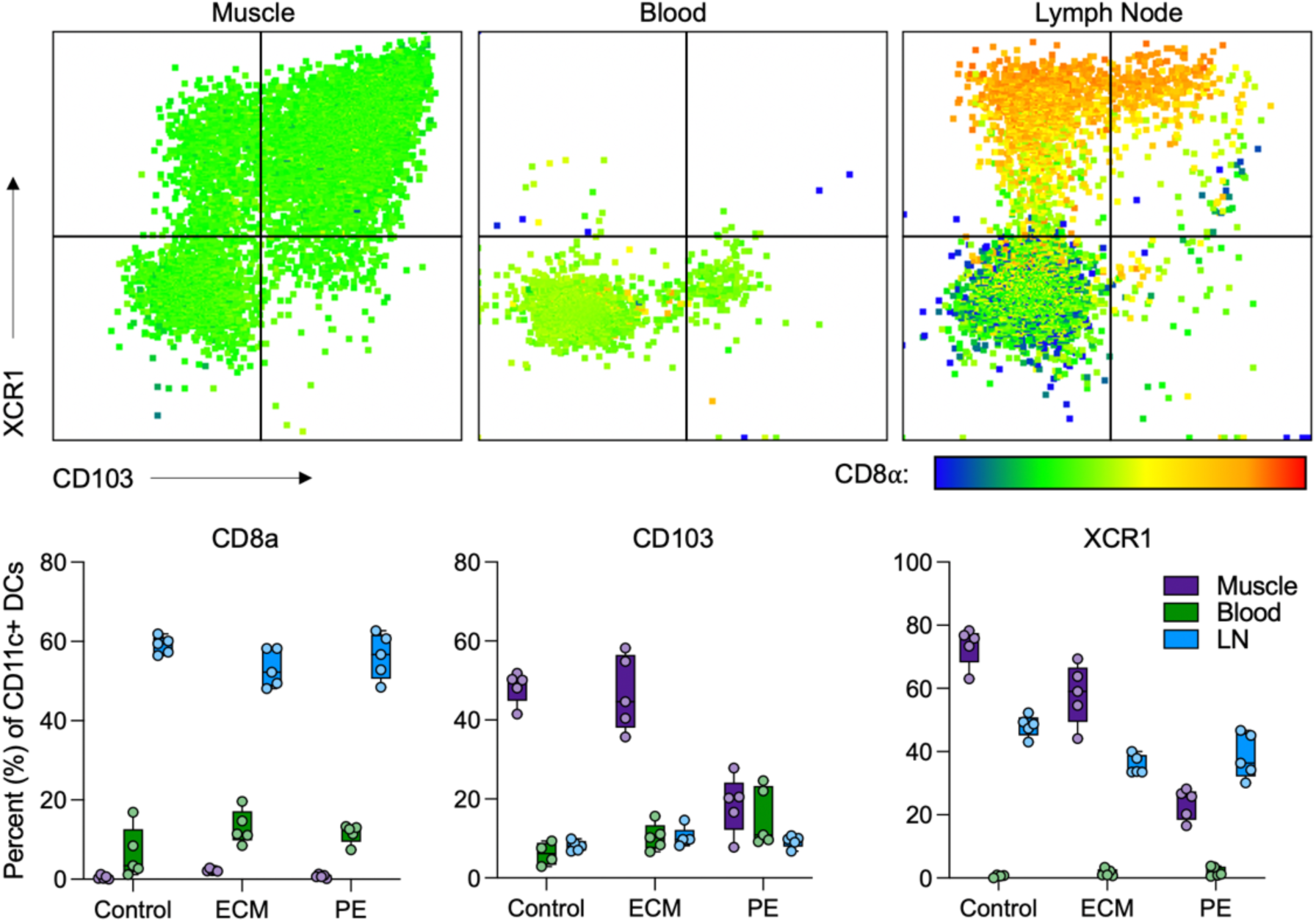
Dendritic cell phenotype in local tissue, blood, and draining lymph node. Top row = representative FACS plots, color scale = CD8⍺ expression. Bottom row: purple = local muscle tissue, green = peripheral blood, blue = inguinal lymph node (LN).

**Supplemental Figure 13.**
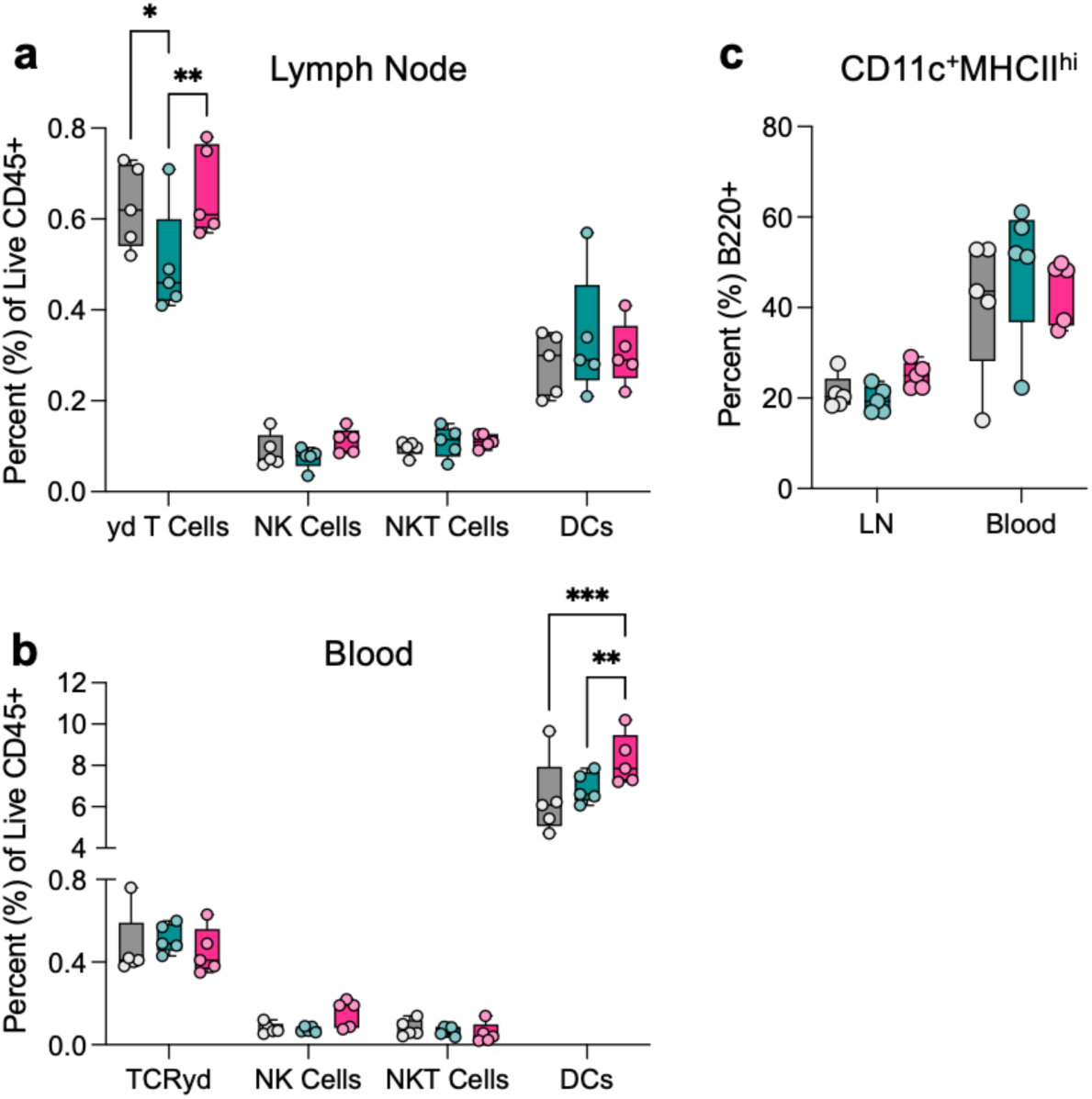
Peripheral plasmacytoid DCs are enriched with pro-fibrotic material treatment of local injury. (a) Innate-like cells in draining lymph node. (b) innate like cells in blood. (c) B220/CD45R expression on dendritic cells in the blood and LN. * = *P* < 0.05; ** = *P* < 0.01, *** = *P* < 0.001. *n* = 5, ANOVA with Tukey post-hoc.

**Supplementary Figure 14.**
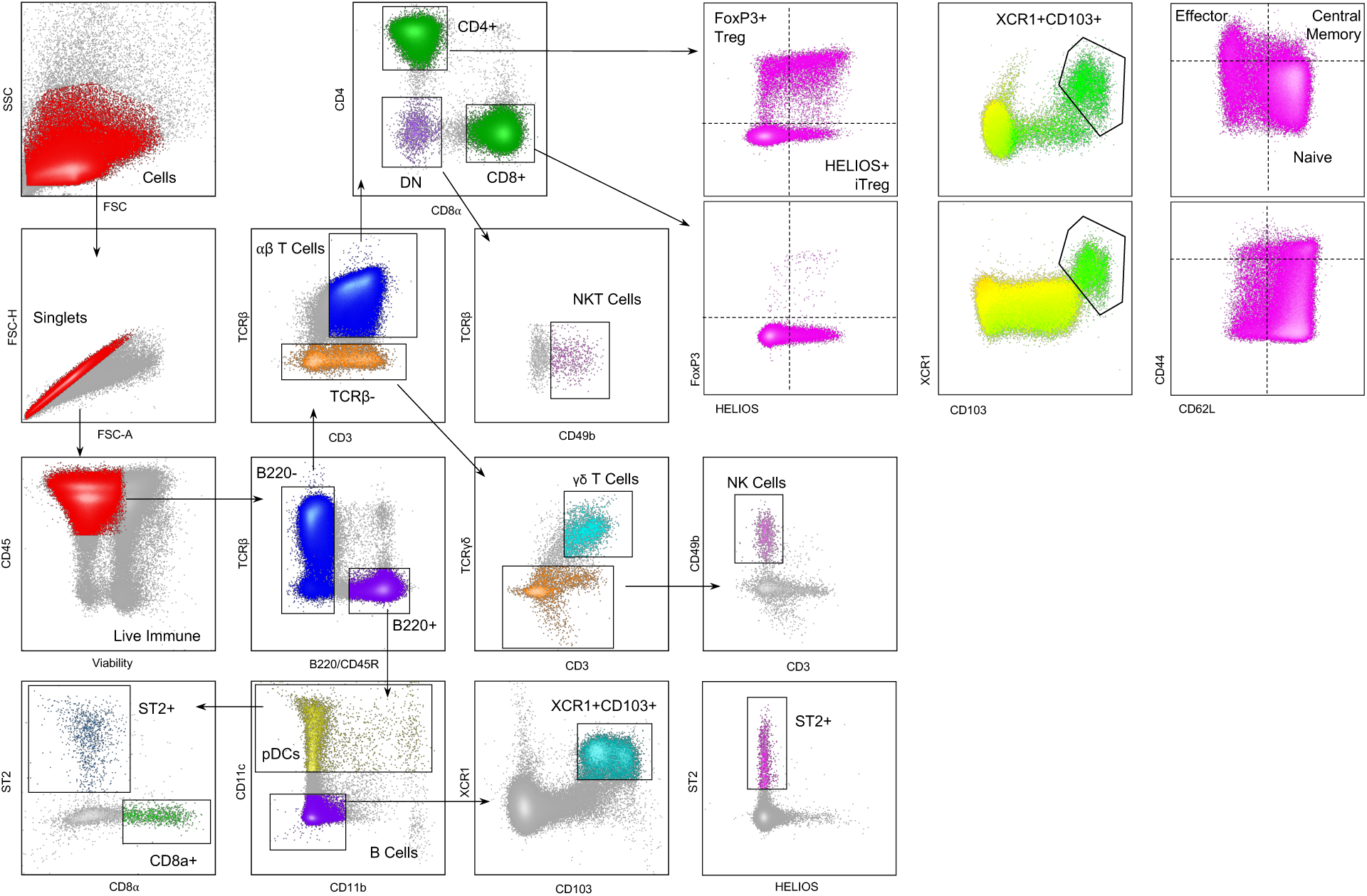
ST2 expressing regulatory B cells are upregulated by ECMtx and dependent upon BATF3. ST2^+^ B cells as a proportion of total B Cells (CD11b^-^CD11c^-^CD49b^-^TCRβ^-^B220^+^). Data are means ± SD, *n* = 3 – 5. ANOVA with Tukey post-hoc correction for multiple comparisons. †† = *P* < 0.01, *** = *P* < 0.001.

**Supplemental Figure 15.**
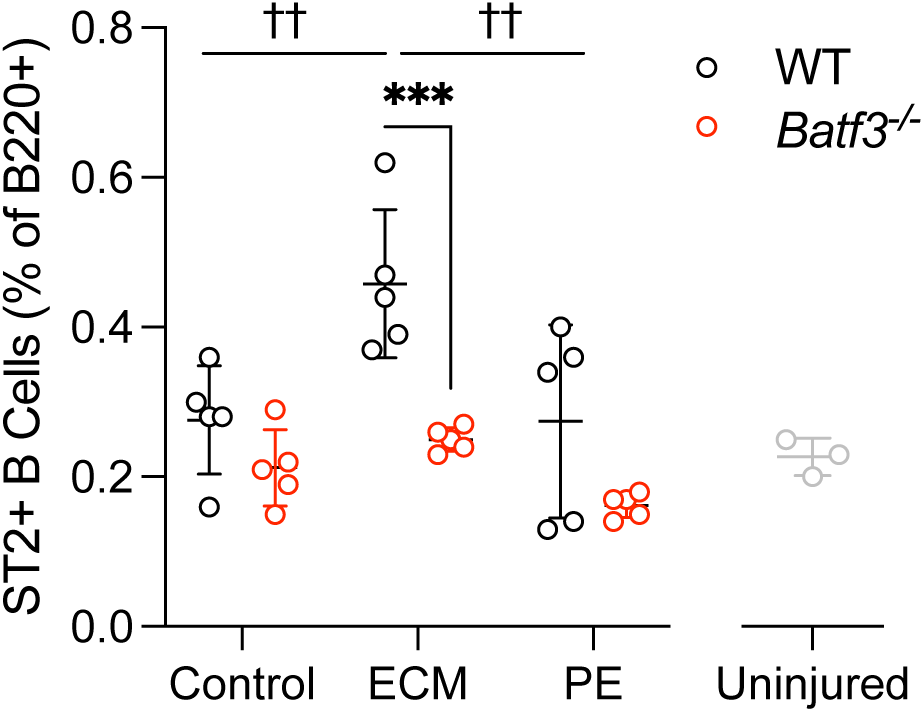
Expansion of active CD103^+^XCR1^+^ adaptive immune cell populations in the draining lymph node after injury. Cells isolated from inguinal lymph nodes at 21 days post-injury. Top panel = proportion of cells that are CD103^+^XCR1^+^, Bottom panel = proportion of CD103^+^XCR1^+^ adaptive immune cells that are CD62L^-^ (active). Uninjured = grey Ⓧ, Control injury = black . *n* = 3 (uninjured), *n* = 5 (control injury). All comparisons (Uninjured vs control injury) are significantly different, *P* < 0.05, ANOVA with Tukey Post-hoc correction for multiple comparisons.

**Supplemental Figure 16.**
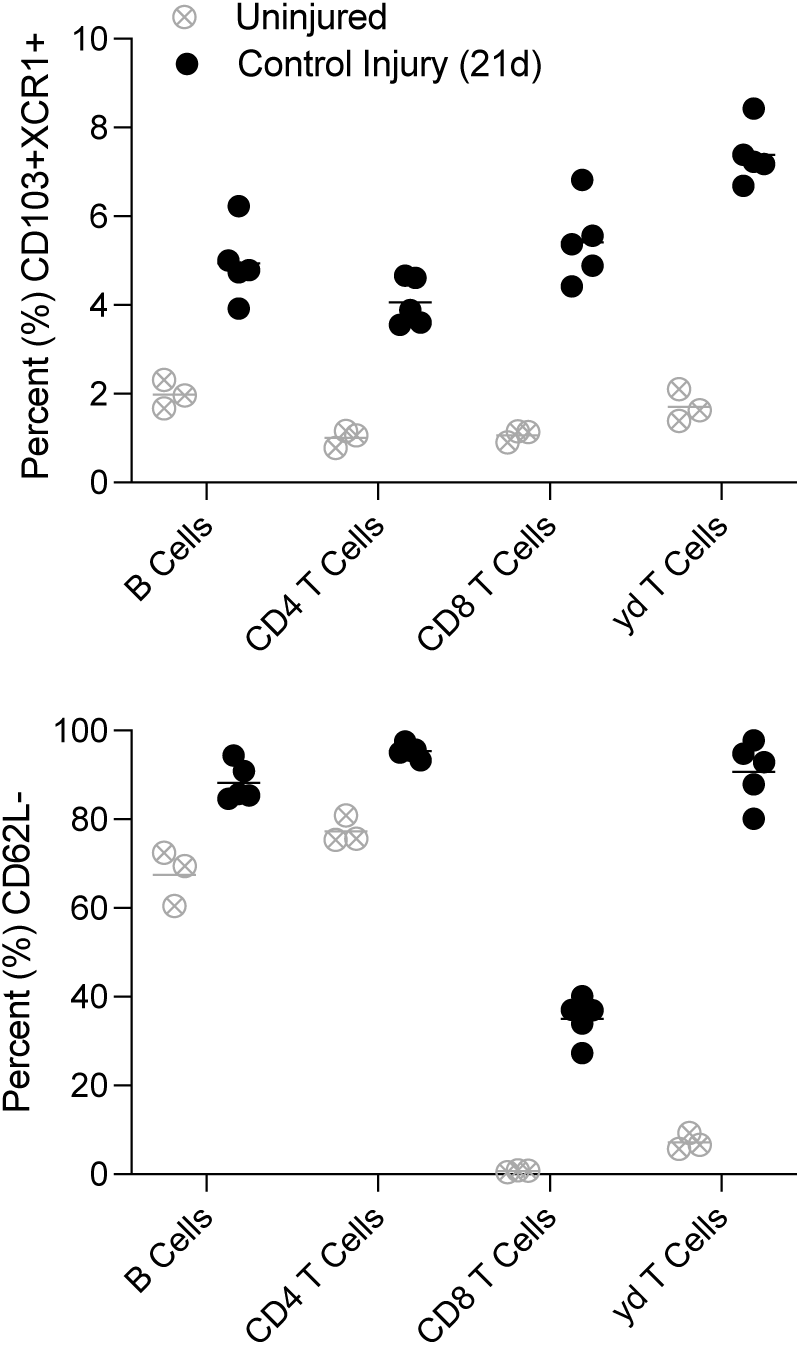
Upregulation and temporal patterning of CD103^+^XCR1^+^ adaptive immune cells after trauma and material implantation. (a) Representative dot plot from draining (inguinal) lymph node from an uninjured mouse and those from the DLN of an injured mouse at 21 days post-injury. B cells = CD45^+^CD11c^-^CD11b^-^CD3^-^B220^+^, T cells = CD45^+^B220^-^CD11c^-^TCRβ^+^CD3^+^CD49b^-^ CD4^+^ (CD4^+^ T Cells) or CD8^+^ (CD8^+^ T cells), γδ T cells = CD45^+^B220^-^TCRβ^-^CD49b^-^TCRγδ^+^. (b) Proportion of adaptive immune cells that are CD103^+^XCR1^+^ (c) Proportion of CD103^+^XCR1^+^ adaptive immune cells that are CD62L^-^ (d) Proportion of adaptive immune cells that are CD103^lo^XCR1^-^ (e) Proportion of CD103^lo^XCR1- adaptive immune cells that are CD62L^-^. Control = black, ECM = teal, PE = pink. Data are means ± 95% Confidence Interval, *n* = 3 – 5.

**Supplemental Figure 17.**
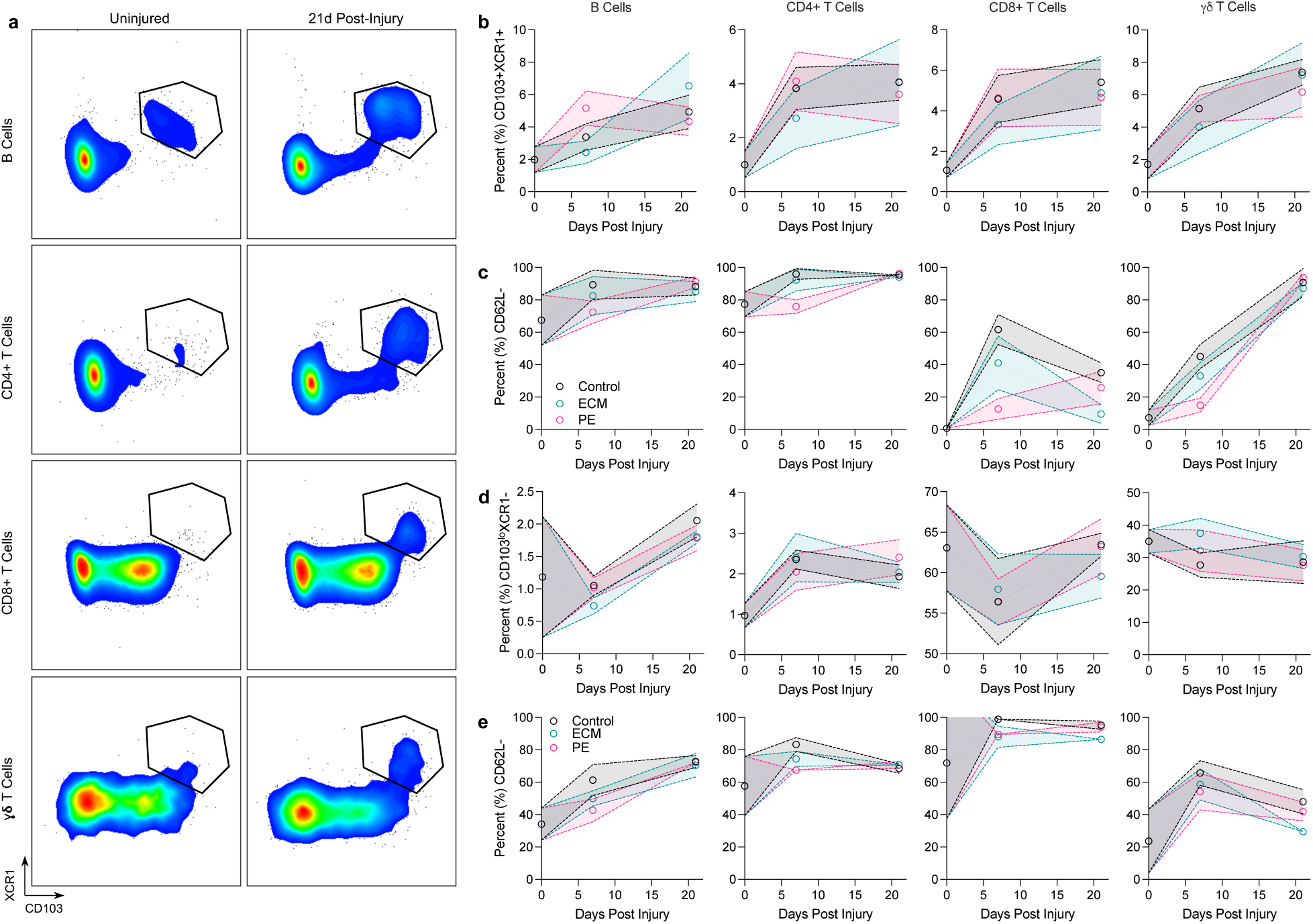
Comparison of CD103^+^XCR1^+^ adaptive immune cells in draining lymph node and peripheral blood at 7- and 21-days post injury. Proportion of adaptive immune cells that are positive for both XCR1 and CD103 in the draining (inguinal) lymph node (ILN) and peripheral blood. Data are means ± SEM, *n* = 4 – 5 αβ T Cells

**Supplemental Figure 18.**
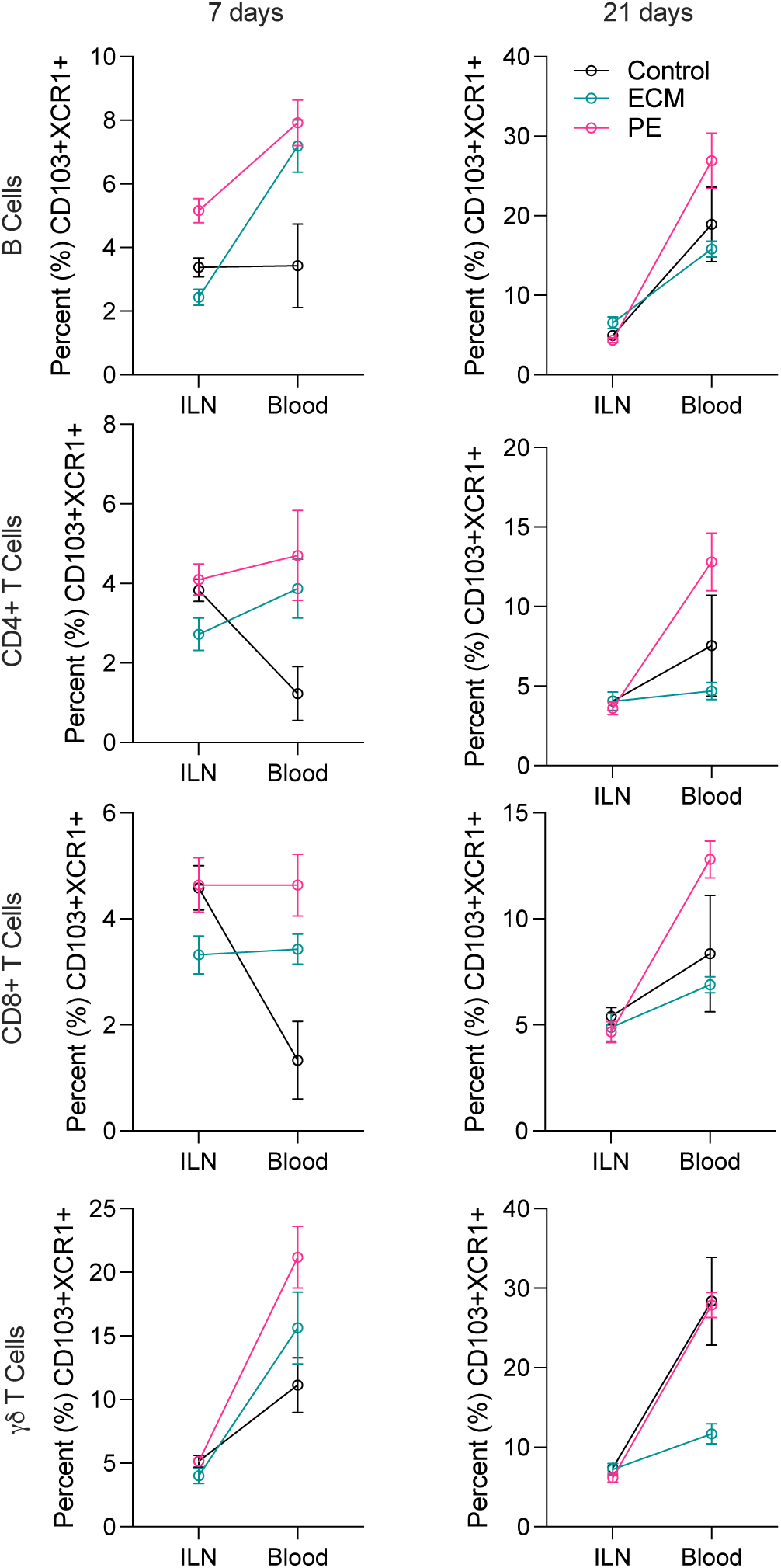
CD103^+^XCR1^+^ T Cells are active and enriched in local tissue compared to draining lymph node. Representative FACS plot showing CD62L and CD44 expression on CD4^+^ T cells that are double positive for CD103 and XCR1 (blue) or double negative (grey) (left). Proportion of CD103^+^XCR1^+^ T cells in the draining lymph node (inguinal), peripheral blood, and muscle injury site (right). Data are means ± SEM, *n* = 5.

**Supplemental Figure 19.**
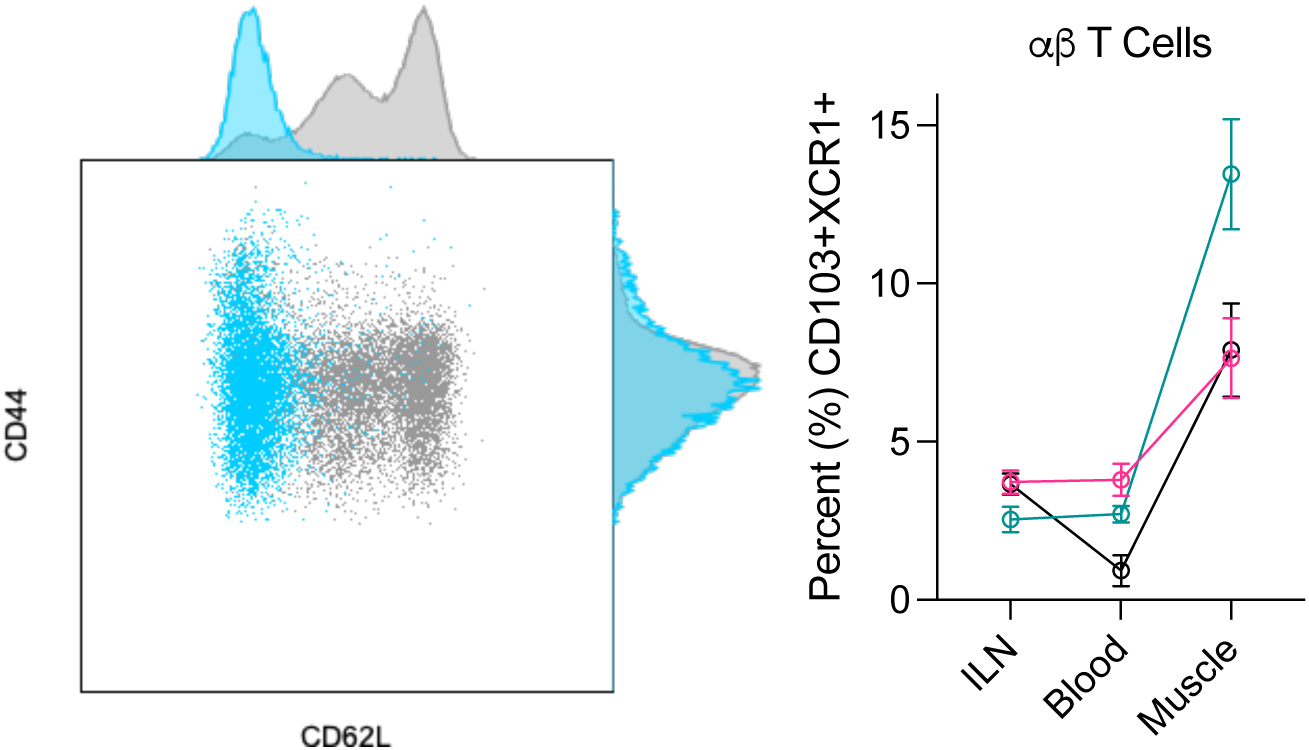
CD103 and XCR1 expression on FoxP3^+^ and HELIOS^+^ regulatory CD4 and CD8 T cells over time. Black = control injury, teal = ECM treated, pink = PE treated. Data are means ± SEM, *n* = 5.

**Supplemental Figure 20.**
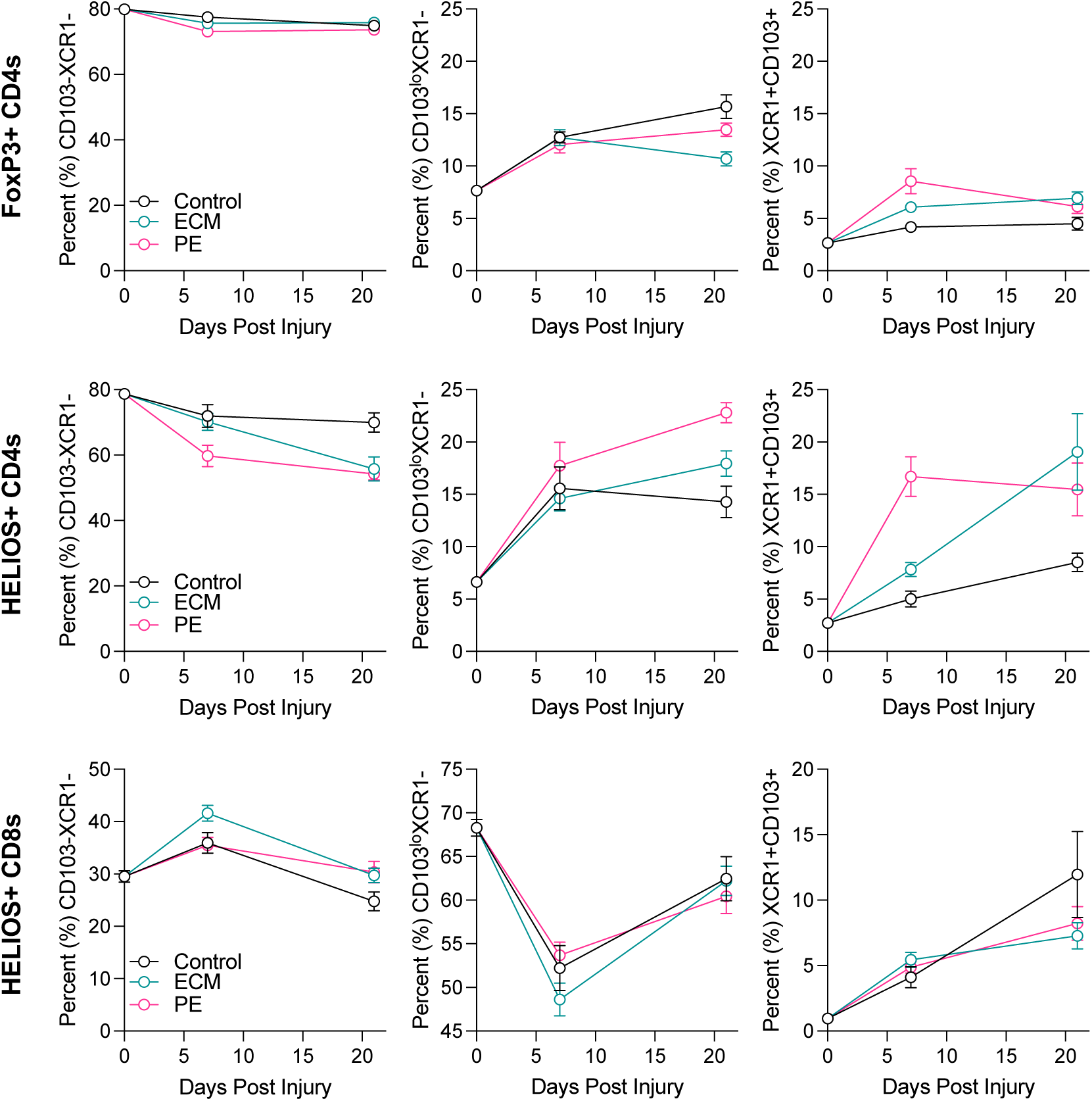
B Cell and γδ T Cell phenotyping in WT and *Batf3^-/-^* mice. Proportion of (a) B cells and (b) γδ T cells that are CD103^+^XCR1^+^ in WT (black) and *Batf3^-/-^* (red) mice in the inguinal (draining) lymph node. (c) B cell, ⍺β T cell, and γδ T cell prevalence in draining lymph node as proportion (%) of CD45^+^ cells. (d) γδ T cell activation in the draining lymph node at 7 days post injury and peripheral blood at 21 days post injury. Data are means ± SD (a-b) or range (c-d), *n* = 3 – 5.

**Supplemental Figure 21.**
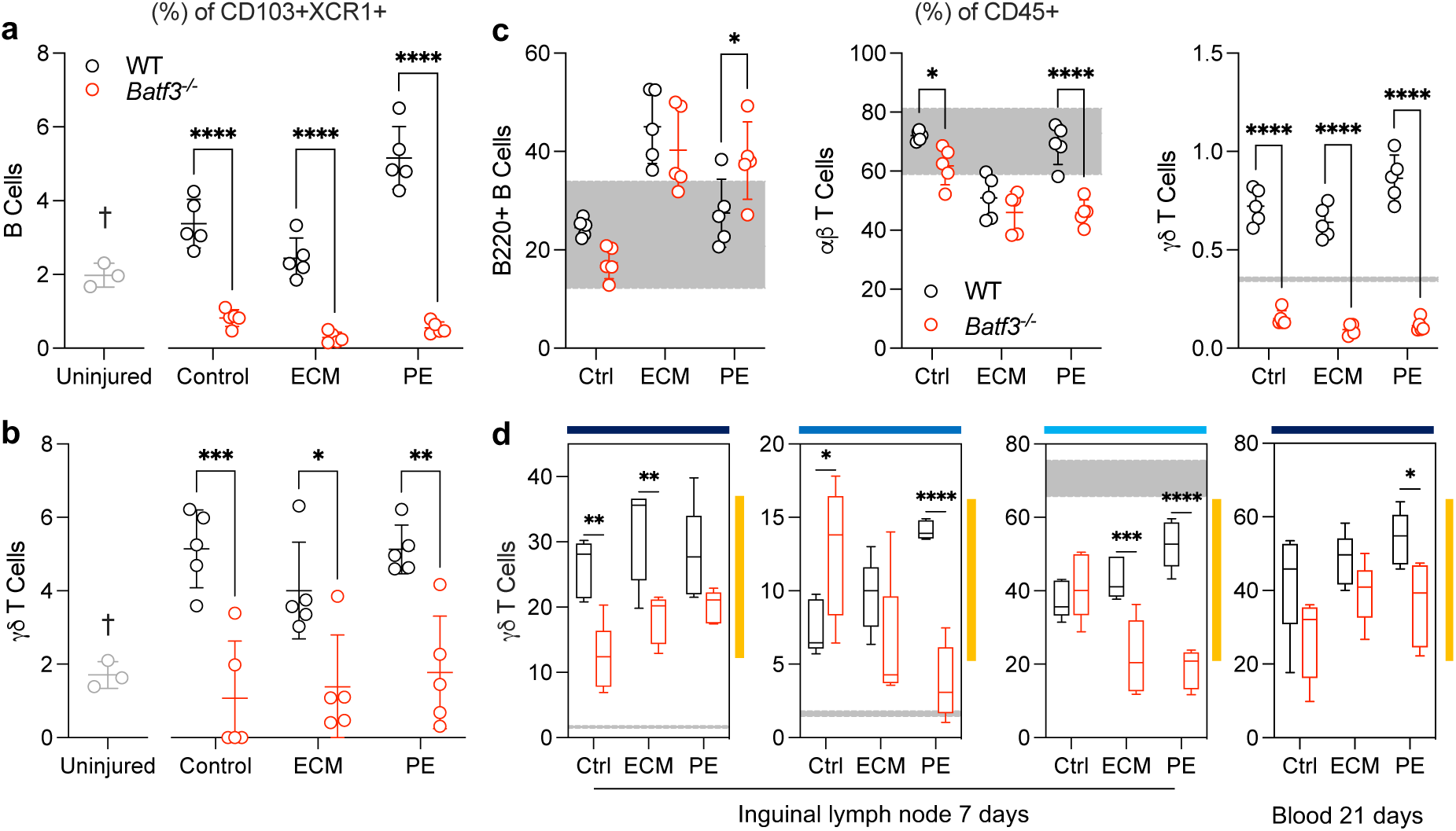
XCL-1 Plasma levels at 7 days post injury. Blue = WT, Red = *Rag1^-/-^*. Data are min to max, *n* = 5. ANOVA with Tukey’s post-hoc correction for multiple comparisons.

**Supplemental Figure 22.**
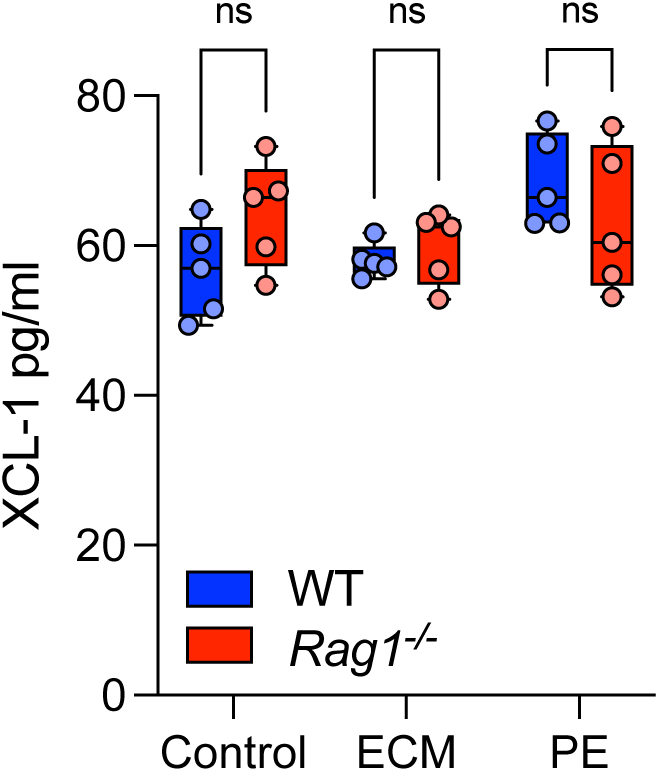
Cytokine and Chemokine profile of local muscle tissue at 7 days post injury. Displayed as a fold change over uninjured control, Control (injury) = Black, ECM = teal, PE = pink. Representative of *n* = 5 blots.

**Supplemental Figure 23.**
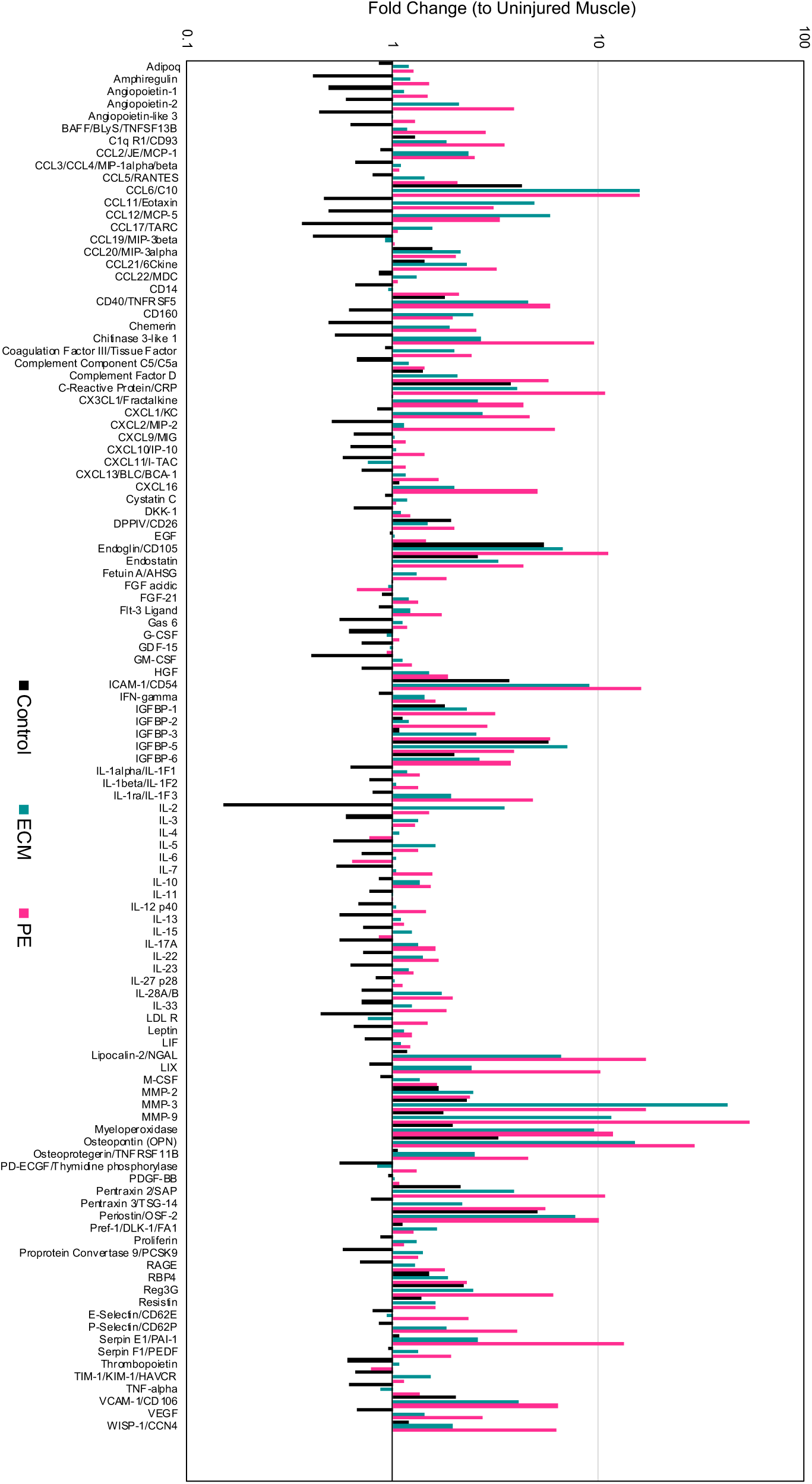
Proposed mechanisms of tissue homeostasis and damage response through communication between CD103+XCR1+ innate and adaptive immune cells. Figure made in BioRender.

**Supplemental Figure 24.**
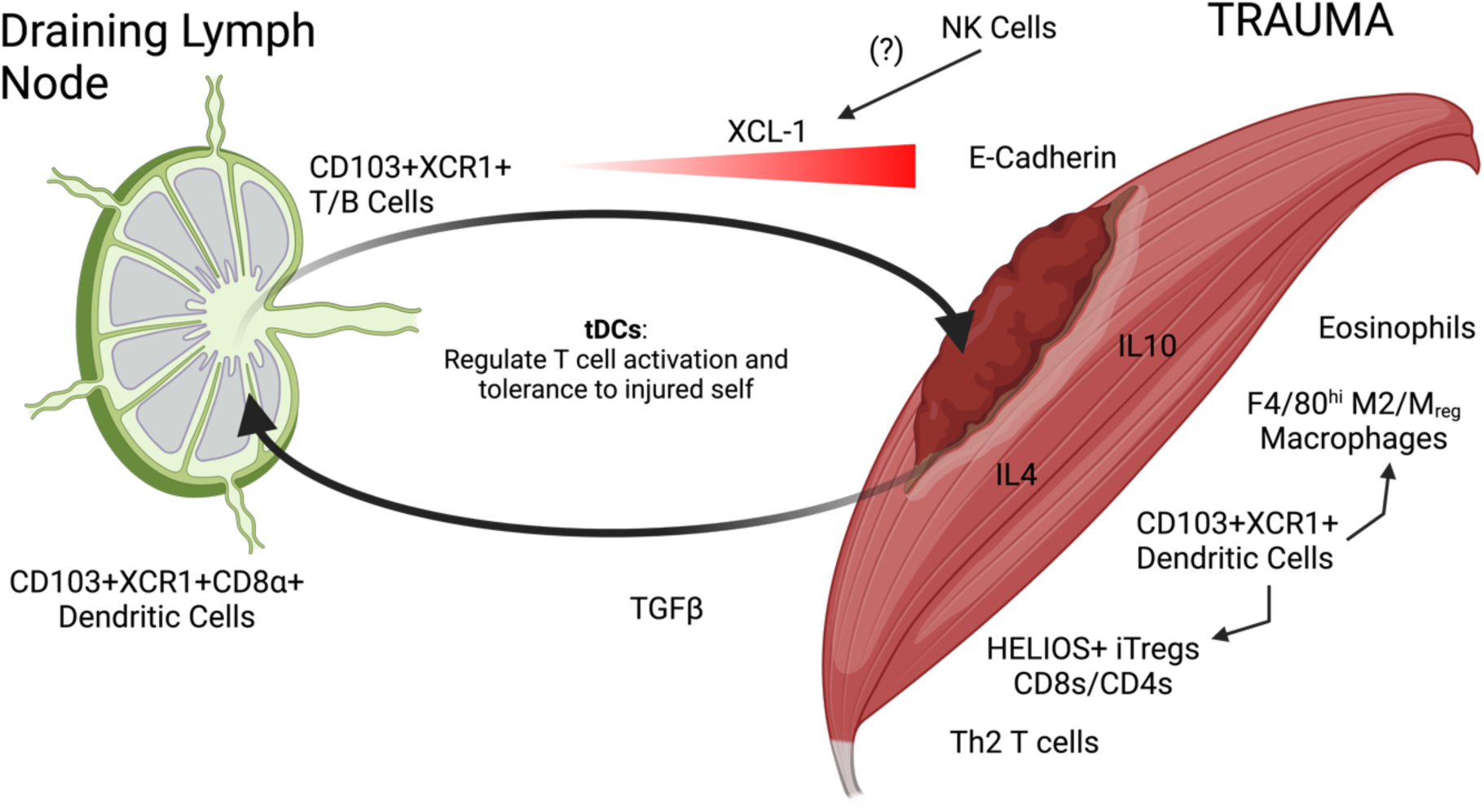
Gating strategy for myeloid panel. Representative plots and gates from sample stained with 22 color myeloid phenotyping panel. Example data are from 7 days post-injury.

**Supplemental Figure 25.**
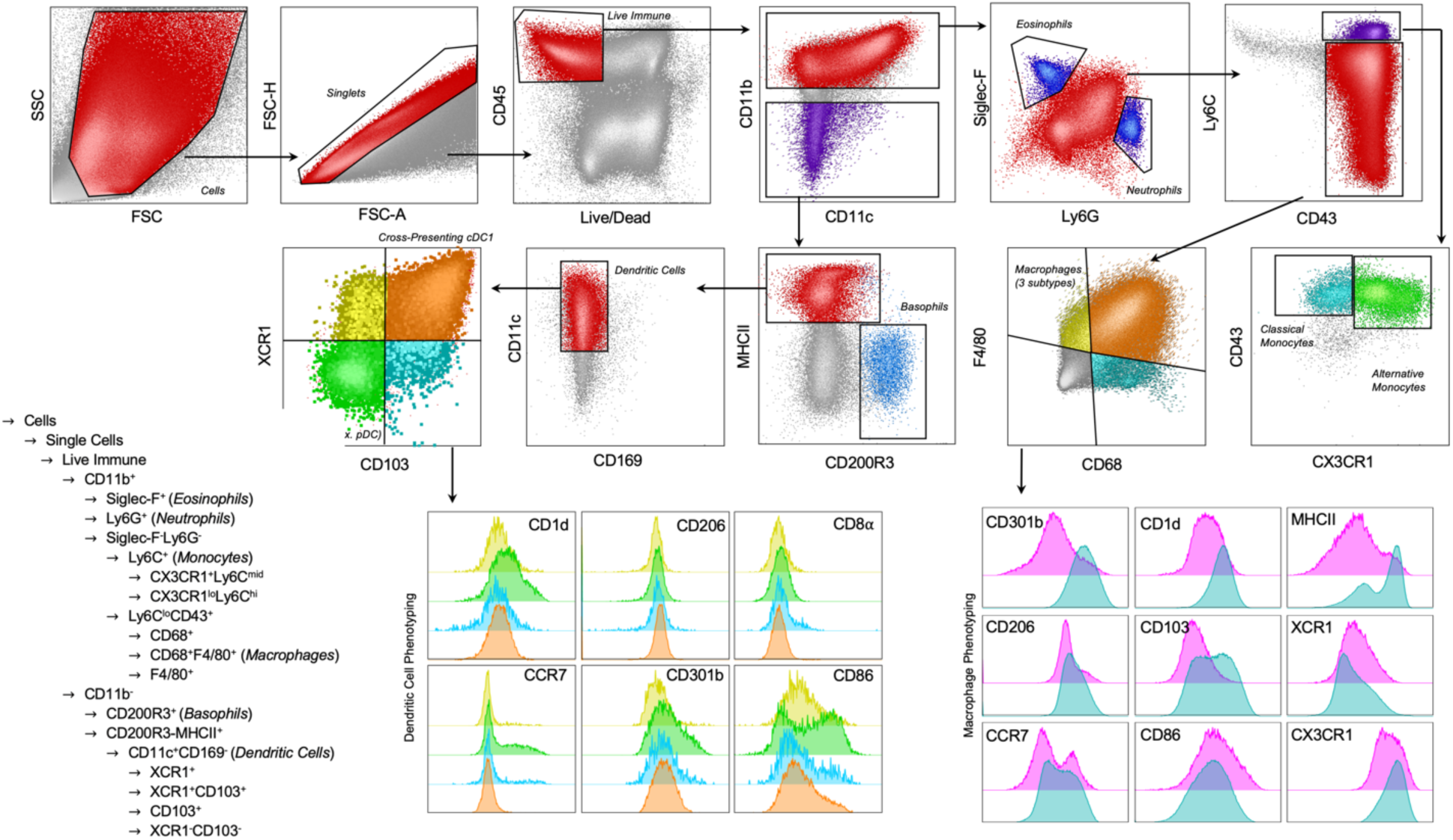
Fluorescence Minus One (FMO) Controls for select myeloid markers. FMO’s from select myeloid panel markers at 7 days post-injury in macrophage and dendritic cell populations. Grey = unstained control, black dashed line = FMO, red = full stained control.

**Supplemental Figure 26.**
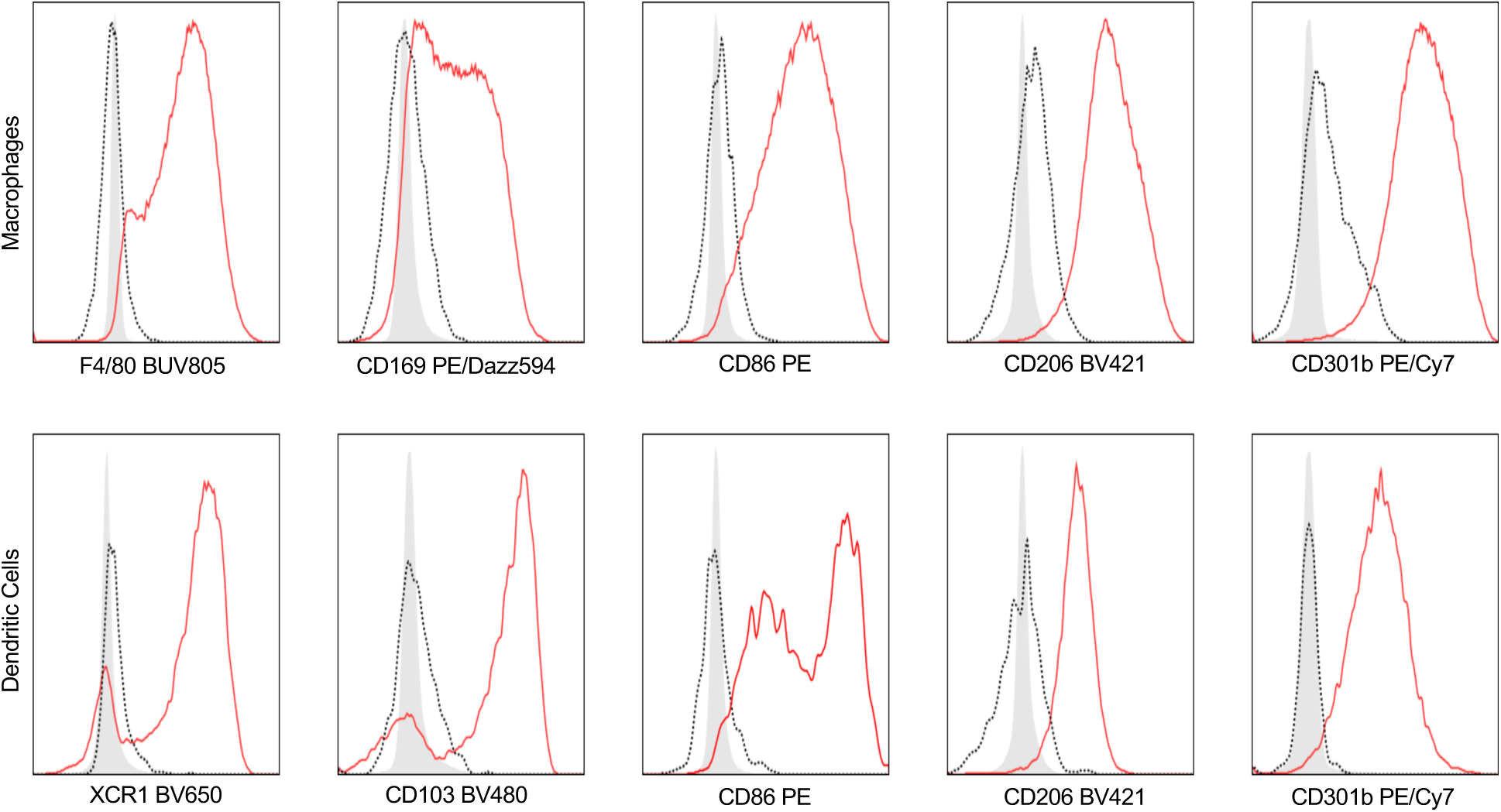
Lymphoid panel gating strategy. Displayed is a representative sample from the inguinal lymph node at 21 days post-injury.

**Supplemental Table 1.** Antibody panel for myeloid phenotyping. Marker = protein antigen, Target = cell population of interest, Dilution Factor = the final dilution of the antibody within the staining cocktail, Bead or Cell Control = the type of sample used for single color control for the spectral unmixing.

| Marker | Target | Clone | Fluorophore | Vendor | Dilution Factor | Bead or Cell Control |
| --- | --- | --- | --- | --- | --- | --- |
| CD45 | Leukocytes | 30-F11 | BUV395 | BD | 1:100 | PBMC |
| Viability | Dead Cells |  | LIVE DEAD Blue | Thermo | 1:1000 | RAW264.7 |
| MHCII | APC | 2G9 | BUV496 | BD | 1:400 | PBMC |
| CD1d | Lipid Ag Present | 1B1 | BUV563 | BD | 1:200 | Beads |
| F4/80 | Macrophage | T45-2342 | BUV805 | BD | 1:100 | Beads |
| CD206 | M2 Mac | C068C2 | BV421 | Biolegend | 1:100 | Beads |
| Ly6G | Neutrophil | 1A8 | eFluor 450 | Thermo | 1:200 | Beads |
| CD103 | DC | M290 | BV480 | BD | 1:200 | Beads |
| Ly6C | Monocyte | HK1.4 | BV510 | Biolegend | 1:200 | PBMC |
| CD11c | DC | N418 | BV570 | Biolegend | 1:200 | Beads |
| Siglec-F | Eosinophil | E50-2440 | BV605 | BD | 1:200 | Beads |
| XCR1 | DC | ZET | BV650 | Biolegend | 1:400 | Beads |
| CD43 | Monocyte | S11 | FITC | Biolegend | 1:200 | PBMC |
| CD8α | DC | 53-6.7 | Spark Blue 550 | Biolegend | 1:100 | PBMC |
| CD301b | Macrophage | URA-1 | PE/Cy7 | Biolegend | 1:400 | PBMC |
| CD86 | M1 Mac | A17199A | PE | Biolegend | 1:400 | PBMC |
| CD169 | Tissue Resident | 3D6.112 | PE/Dazzle594 | Biolegend | 1:400 | Beads |
| CCR7 | M1 Mac | 4B12 | PE-Cy5 | Biolegend | 1:200 | PBMC |
| CD200R3 | Baso/Mast Cell | Ba13 | APC | Biolegend | 1:200 | Beads |
| CX3CR1 | Pre-M2 | SA011F11 | Alexa Fluor 647 | Biolegend | 1:400 | PBMC |
| CD11b | Myeloid | M1/70 | Alexa Fluor 700 | Biolegend | 1:400 | PBMC |
| CD68 | Macrophage | FA-11 | APC-Cy7 | Biolegend | 1:100 | Beads |

**Supplemental Table 2.**
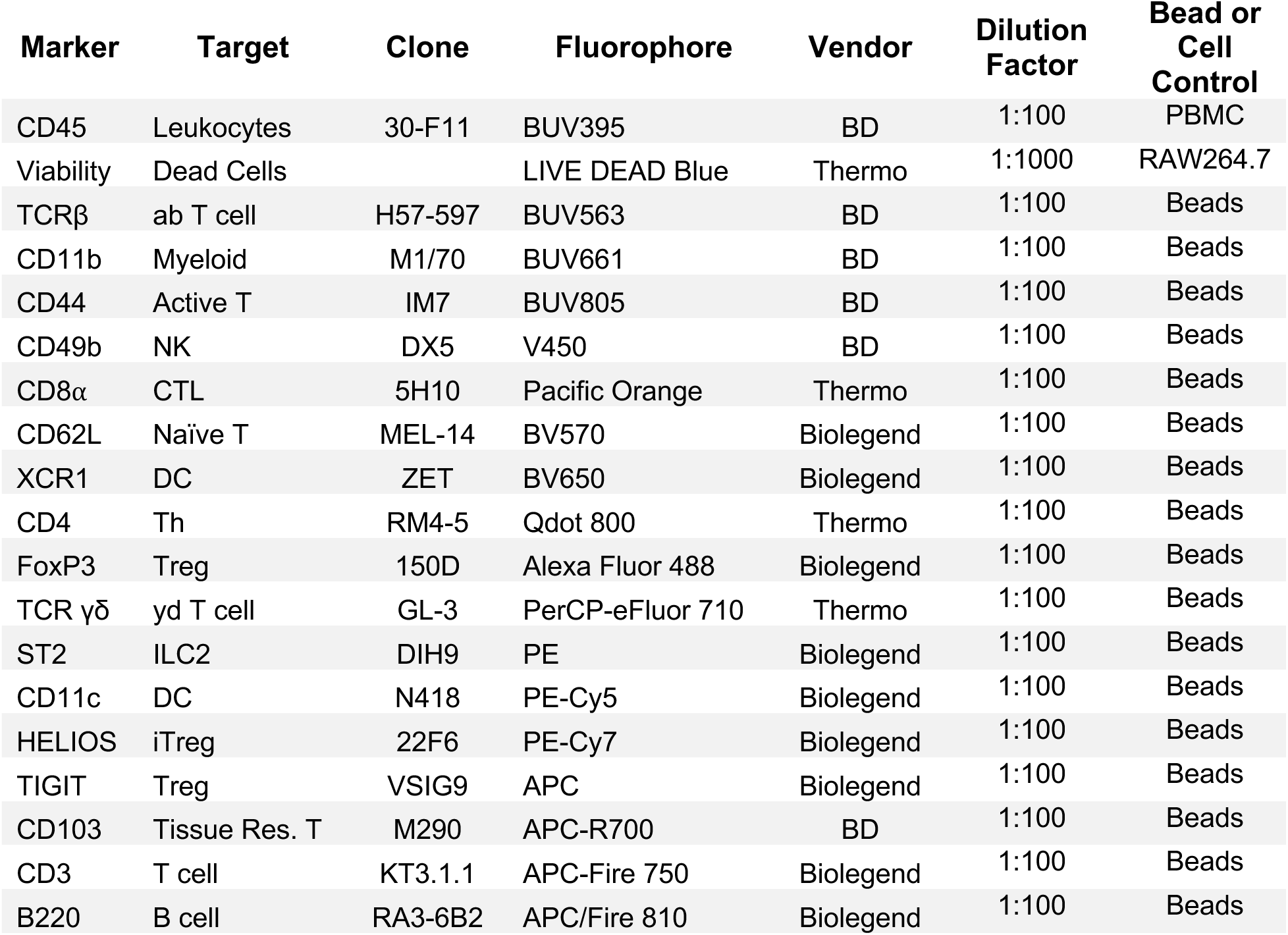
Antibody Panel for lymphoid phenotyping. Marker = protein antigen, Target = cell population of interest, Dilution Factor = the final dilution of the antibody within the staining cocktail, Bead or Cell Control = the type of sample used for single color control for the spectral unmixing.

**Supplementary Table 3.** TaqMan RT-PCR Primer Probes. Gene, reporter, and probe ID are listed. Housekeeping genes used in normalization are denoted with an asterix (*). Probes were selected to span exon-exon junctions.

| Gene | Reporter | Probe ID |
| --- | --- | --- |
| *Gusb* * | FAM-MGB | Mm01197698_m1 |
| *Xcl1* | FAM-MGB | Mm00434772_m1 |
| *Il10* | FAM-MGB | Mm01288386_m1 |
| *Il4* | FAM-MGB | Mm00445259_m1 |
| *Cdh1* | FAM-MGB | Mm01247357_m1 |
| *Tgfb1* | FAM-MGB | Mm01178820_m1 |
| *Xcr1* | FAM-MGB | Mm00627650_m1 |

